# Modeling of Alpha-1 Antitrypsin Deficiency with Syngeneic Human iPSC-Hepatocytes Reveals Metabolic Dysregulation and Cellular Heterogeneity in PiMZ and PiZZ Hepatocytes

**DOI:** 10.1101/2022.02.01.478663

**Authors:** Joseph E Kaserman, Rhiannon B. Werder, Feiya Wang, Taylor Matte, Michelle I. Higgins, Mark Dodge, Jonathan Lindstrom-Vautrin, Anne Hinds, Esther Bullitt, Ignacio S. Caballero, Xu Shi, Robert E. Gerszten, Nicola Brunetti-Pierri, Marc Liesa, Carlos Villacorta-Martin, Anthony N. Hollenberg, Darrell N. Kotton, Andrew A. Wilson

## Abstract

Individuals homozygous for the pathogenic “Z” mutation in alpha-1 antitrypsin deficiency (AATD) are known to be at increased risk for chronic liver disease. That some degree of risk is similarly conferred by the heterozygous state, estimated to affect 2% of the US population, has also become clear. A lack of model systems that recapitulate heterozygosity in human hepatocytes has limited the ability to study the impact of expressing a single ZAAT allele on hepatocyte biology. Here, through the application of CRISPR-Cas9 editing, we describe the derivation of syngeneic induced pluripotent stem cells (iPSCs) engineered to determine the effects of ZAAT heterozygosity in iPSC-derived hepatocytes (iHeps) relative to homozygous mutant (ZZ) or corrected (MM) cells. We find that heterozygous MZ iHeps exhibit an intermediate disease phenotype and share with ZZ iHeps alterations in AAT protein processing and downstream perturbations in hepatic metabolic function including ER and mitochondrial morphology, reduced mitochondrial respiration, and branch-specific activation of the unfolded protein response in subpopulations of cells. Our cellular model of MZ heterozygosity thus provides evidence that expression of a single Z allele is sufficient to disrupt hepatocyte homeostatic function and suggest a mechanism underlying the increased risk of liver disease observed among MZ individuals.

## Introduction

Alpha-1 antitrypsin deficiency (AATD) is a common inherited cause of chronic liver disease, driven by accumulation of misfolded protein aggregates and associated deleterious effects (1–3). The most common disease-causing variant, known as the “Z” mutation, is a point mutation within the *SERPINA1* gene resulting in an amino acid substitution (Glu342Lys), that predisposes nascent Z alpha-1 antitrypsin (ZAAT) proteins to misfold and polymerize within the endoplasmic reticulum (ER) of hepatocytes (4–6). It has long been recognized that “PiZZ” homozygous individuals (hereafter referred to as ZZ) are at increased risk for chronic liver disease (4–6). Whether some degree of increased risk is associated with the heterozygous state has only recently become clear. While older studies generated conflicting results, more recent evidence has identified a modest increased risk for clinically significant liver disease among heterozygous “PiMZ” individuals (hereafter referred to as MZ), particularly in the context of a second injury (7–13).

The lack of a model system that faithfully reproduces human MZ hepatocyte biology hindered the determination of whether ZAAT heterozygosity can induce injury in human hepatocytes as well as a direct comparison to the better-characterized effect of ZAAT homozygosity. Transgenic “PiZ” mice co-express murine AAT together with multiple copies of the human Z allele (14,15), while immortalized cell lines engineered to express Z- or M-AAT (16,17) fail to capture patient-to-patient genetic heterogeneity or hepatocyte-specific biology that could contribute to variable risk among individuals. Given that MZ heterozygotes represent approximately 2% of the US population, there is a critical need to establish models that capture genetic diversity and are likewise capable of recapitulating heterozygosity in human liver cells (1,18).

To complement existing models, we and others have applied ZZ patient-specific induced pluripotent stem cell (iPSC)-derived hepatocytes (iHeps) to recapitulate cellular features of ZAAT-associated liver disease pathogenesis (19–24). Here, we extend this work to directly test the impact of ZAAT heterozygosity on hepatocyte biology using genetically controlled syngeneic MZ and MM iHeps generated from three distinct ZZ patient-specific iPSC lines. Through a combination of bulk and single cell RNA sequencing and metabolomics analysis, we find that MZ iHeps exhibit an intermediate phenotype and share with ZZ iHeps significant alterations in AAT protein processing associated with downstream metabolic dysregulation and branch-specific activation of the unfolded protein response (UPR) among cellular subsets.

## Results

### Generation of Z mutant heterozygous human iPSCs

To test the hypothesis that expression of a single Z allele is sufficient to promote liver injury, we decided to repair the Z mutation in iPSCs derived from highly phenotyped ZZ homozygous patients (19,24). To do so, we used CRISPR/Cas9 in combination with a single stranded oligodeoxynucleotide (ssODN) donor template, utilizing a protocol we have previously applied for efficient correction of the Z mutation (19). To correct one or both alleles, we introduced either one or two ssODN donors containing the Z or wild type (M) sequence (**Figure 1A**) (25). Utilizing this approach, we successfully targeted 3 genetically distinct ZZ iPSC lines (referred to as PiZZ1, PiZZ6, and PiZZ100) generating 3 syngeneic sets of ZAAT homozygous (ZZ), heterozygous (MZ), and wild type (MM) iPSCs for a total of 9 lines (**Figures 1A, S1A**). Following targeting, we confirmed that daughter iPSC lines retained a normal karyotype (**Figure S1B**) and found no evidence of off target editing through sequencing of the 6 top computationally-predicted potential off target genomic sites in each line (data not shown).

**Figure 1:**
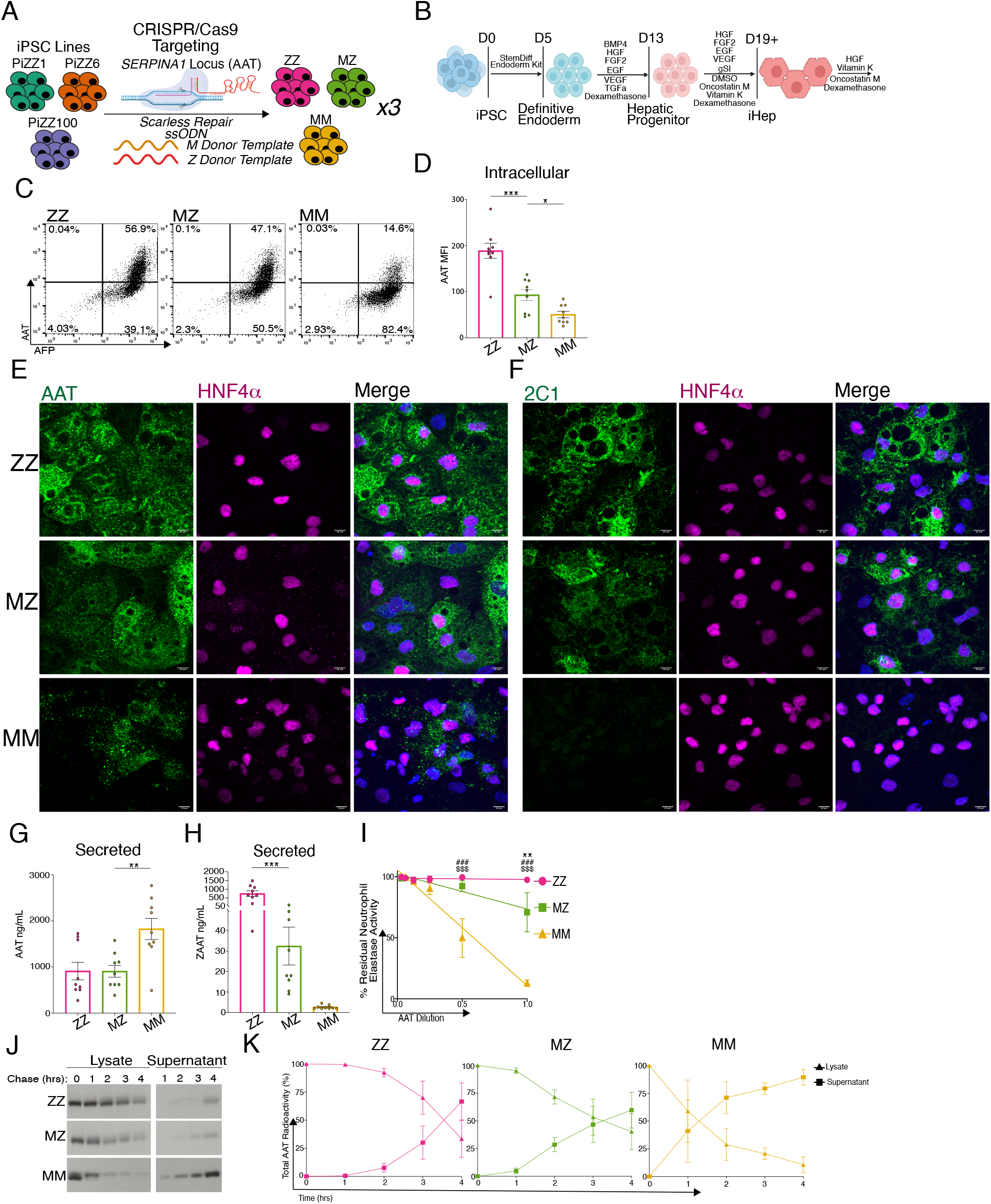
Characterization of Syngeneic MZ and MM iHeps from CRISPR/Cas9 Edited ZZ iPSCs. A) Targeting strategy for the *SERPINA1* (AAT) locus. B) Schematic of directed differentiation protocol for generating iHeps. C) Representative flow cytometry plots of fixed, permeabilized ZZ, MZ, and MM iHeps. D) MFI of intracellular AAT protein in ZZ, MZ and MM iHeps. E, F) Immunostaining of ZZ, MZ and MM iHeps for AAT, 2C1 and HNF4α. G) Secreted total AAT and ZAAT (H) protein in ZZ, MZ and MM iHep supernatants. I) Assay of anti-neutrophil elastase inhibition in concentrated iHep supernatants. J) Representative quantification of AAT secretion kinetics using ^35^S-Met/Cys labeled AAT protein from intracellular protein lysates and supernatants. K) Kinetic of aggregated AAT labeled cell lysate and supernatants from (J). n = 3 independent experiments from each of the syngeneic backgrounds. Data represented as mean +/- SEM. *p<0.05, **p<0.01, ***p<0.001 by one-way anova with Dunnett’s multiple comparisons test using MZ as control. Neutrophil elastase inhibition ZZ vs MZ **p<0.01, ZZ vs MM ^###^p<0.001, MZ vs MM ^$$$^p<0.001 by two-way anova with Tukey’s multiple comparisons test.

### A single Z allele is sufficient to alter iHep intracellular AAT trafficking

We next sought to characterize AAT intracellular protein processing in MZ iHeps relative to genetically matched ZZ and MM iHep comparators (n=3 genetically distinct iPSC lines per group). To derive iHeps from undifferentiated iPSCs, we applied a directed differentiation protocol we have previously shown to efficiently generate hepatic cells that transcriptomically resemble primary human hepatocytes and recapitulate critical features of AATD pathobiology (**Figures 1B, S1C**), (19,24). Consistent with the known consequences of protein misfolding associated with mutant ZAAT expression (4,5,24), ZZ iHeps demonstrated significant intracellular AAT protein retention as quantified by flow cytometry (**Figures 1C,D, S1D**). Based on percentage of cells staining positive for AAT and on the mean fluorescence intensity (MFI) of antibody staining, intracellular AAT protein levels were lower in MZ iHeps than in ZZ iHeps, but significantly higher than in MM iHeps (**Figures 1C,D, S1D**). To determine whether intracellular AAT formed polymers, we performed immunostaining using either a pan-AAT antibody or the 2C1 antibody that specifically recognizes AAT polymers (**Figures 1E,F, S1E**) (26). Similar to the flow cytometric results, pan-AAT antibody staining correlated with the number of iHep Z alleles, generating the least signal in MM cells. Staining with the 2C1 antibody was positive in MZ and ZZ, but not MM, iHeps, indicating the presence of polymerized AAT in homozygous mutant and heterozygous cells. Next, we quantified secreted AAT by ELISA, again using a pan-AAT antibody or an antibody specific for ZAAT protein (**Figure 1G,H**) (27). Total AAT protein levels in MZ iHep supernatants were similar to ZZ levels but significantly lower than those observed in MM supernatants (**Figure 1G**). Assays employing the ZAAT-specific antibody revealed significantly less ZAAT protein in MZ than in ZZ iHep supernatants (**Figures 1H, S1D,F**), suggesting that AAT secreted by MZ iHeps is predominantly MAAT. Additionally, we found that the neutrophil elastase inhibitory capacity of concentrated supernatants from MZ iHeps exceeded that of ZZ iHep supernatants (**Figure 1I**).

We next performed pulse-chase ^35^S-Met/Cys radiolabeling to characterize the processing and secretion kinetics of nascent AAT proteins. As expected based on prior observations (24), wild type AAT protein produced by MM iHeps was rapidly glycosylated to produce the mature 55 kDa AAT isoform and secreted into the supernatant, while protein from ZZ iHeps exhibited a delay in both post-translational processing and secretion (**Figure 1J,K**). In contrast, newly synthesized AAT protein in MZ iHeps exhibited an intermediate phenotype characterized by prolonged retention relative to AAT produced in MM cells but more rapid secretion when compared with that from ZZ iHeps (**Figure 1J,K**). Together, these data demonstrate that MZ iHeps exhibit aberrant AAT processing, retention, and secretion in comparison to genetically matched MM controls.

### Transcriptomic analysis reveals activation of ER stress and metabolic dysregulation in heterozygous MZ iHeps

We next looked to see how either mono- or bi-allelic expression of mutant ZAAT might affect the iHep global transcriptome. Using bulk RNA sequencing, we profiled the transcriptome across syngeneic ZZ, MZ, and MM iHeps derived from the parental line “PiZZ6”. Principle component analysis (PCA) demonstrated three distinct clusters separated by AAT genotype (**Figure 2A**). While MM iHeps separated from the other samples by first principle component variance, MZ and ZZ iHeps segregated only by the second principle component variance, consistent with a lesser degree of transcriptomic variation (**Figure 2A**). Consistent with this finding, the total number of differentially expressed genes (DEG) across three pairwise comparisons revealed more DEG between ZZ (1308) or MZ (736) in comparison to MM iHeps than between each other (66; false discovery rate (FDR) <0.05) (**Figure 2B**). Included in the most up-regulated transcripts for both MZ and ZZ relative to MM iHeps were *IGFBP5*, *SLC15A4*, and *GLT1D1*, which have previously been shown to be involved in hepatocyte response to insulin signaling as well as lipogenesis (28,29), consistent with dysregulated cellular metabolism in *SERPINA1* expressing mutant cells. We then compared the relative expression of transcription factors known to regulate lipid homeostasis and found multiple factors including *HNF1A*, *HNF4A*, *CEBPA*, *PPARA*, *CREB3L3 and RXRA w*ere significantly down-regulated (FDR <0.05) in ZZ iHeps as compared to MM iHeps (**Figure 2C**) (30). Interestingly, we observed a similar but lesser down-regulation of these transcription factors in MZ iHeps with a subset (*CEBPA*, *CREB3L3*, and *RXRA*) also reaching significance (**Figure 2C**). To identify potential pathological processes contributing to the ZAAT-driven transcriptomic changes we next applied gene set enrichment analysis (GSEA) (**Figure 2D,E**). This analysis demonstrated significant enrichment in both MZ and ZZ iHep transcriptomes for genes associated with angiogenesis, apoptosis, TGF-β and epithelial mesenchymal transition (EMT) (FDR <0.1). Conversely, the MM iHep transcriptome was enriched in pathways associated with traditional hepatocyte biology including bile acid metabolism, xenobiotic metabolism, coagulation, and fatty acid metabolism. To contextualize these findings, we compared our data with a previously published dataset that compared the transcriptomes of ZZ to MM iHeps following zinc finger nuclease-mediated biallelic correction of the Z mutation (21). We found significant overlap in the numbers of total genes differentially expressed between ZZ and MM iHeps (1675 vs 1308) and among specific Hallmark gene sets significantly enriched in both datasets (**Figure 2D**) (31). We also detected differences between the two datasets as would be expected based on their differing genetic backgrounds. ZAAT-associated Hallmark enrichment was also consistent with gene sets we have previously identified to be enriched in homozygous and heterozygous Z mutant iHeps when compared to non-diseased MM primary human hepatocytes analyzed by gene set variation analysis (GSVA) (19). We next identified gene ontology (GO) terms enriched by genotype and identified within MM iHeps, as compared with either MZ or ZZ iHeps and found that multiple of the top 20 most significantly enriched terms (FDR <0.05, ranked by NES) were associated with synthetic cellular processes and metabolism (**Table S1**). Given the importance of normal homeostatic mitochondrial and ER function in protein synthesis and regulation of hepatocyte metabolism, we looked to see whether pathways associated with dysfunction of either the ER or mitochondria were enriched in MZ and ZZ iHep transcriptomes. We found that both ZZ and MZ iHeps were enriched (FDR < 0.25) for GO terms such as Positive Regulation of Mitochondrial Outer Membrane Permeability Involved in Apoptotic Signaling and PERK Mediated UPR (**Figure 2F**), suggesting dysfunction of both organelles. Taken together, these data demonstrate that mono-allelic expression of ZAAT is sufficient to perturb the global hepatic transcriptome and dysregulate expression of metabolic transcription factors.

**Figure 2:**
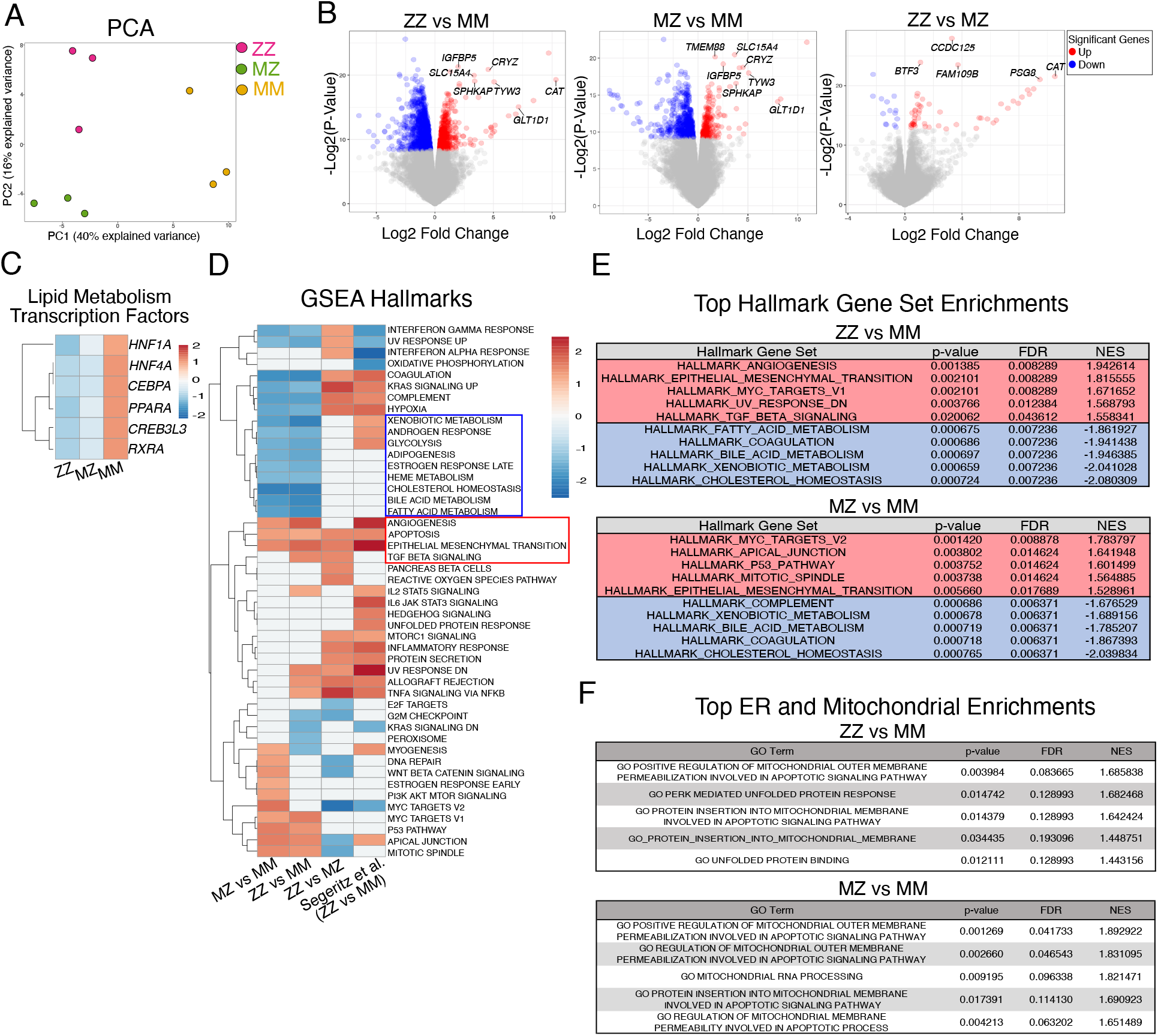
RNA-seq Demonstrates a Single Z Allele is Sufficient to Perturb the Global Transcriptome. A) Principle component analysis (PCA). B) Volcano plots identify differentially expressed genes (DEGs) between ZZ, MZ, and MM iHeps. C) Heatmap of hepatic lipid metabolic transcription factors. D) Gene set enrichment analysis (GSEA) including a published dataset of ZZ vs MM iHeps is included for comparison (FDR <0.1) (21). E) Top 5 most up and down regulated gene sets from ZZ and MZ iHeps as compared to MM iHeps ranked by normalized enrichment score (NES) (FDR <0.05). F) Top 5 ER and mitochondrial associated GO terms ranked by NES (FDR <0.25).

### Metabolomic analyses reveal altered amino acid metabolism, decreased mitochondrial oxidative function and urea cycle defects induced by ZAAT expression

Because multiple metabolic pathways were dysregulated at the transcriptomic level, we next characterized the metabolome to determine which transcriptional changes could be the most relevant to liver metabolism and thus function. To do so, we utilized liquid chromatography-mass spectrometry (LC-MS) with extraction methods to enrich for both lipid and amide metabolites (32). To complement our transcriptomic analysis, we again compared ZZ, MZ, and MM iHeps derived from “PiZZ6” and identified 128 (ZZ vs MM) and 132 (MZ vs MM) total differential metabolites (FDR < 0.05) from the combined lipid and amide fractions (**Table S2**). Branched chain amino acids (BCAA) and short chain acylcarnitines were elevated in both MZ and ZZ relative to MM iHeps while ATP, lactate, reduced glutathione (GSH), oxidized glutathione (GSSG), and long chain acylcarnitines were depleted (**Figure 3A, Table S2**). We also observed a reduction in C2 acylcarnitines in MZ and ZZ iHeps, a pattern suggesting decreased fatty acid oxidation and carnitine shuttle activity within mitochondria (**Figure 3A**). Additionally, multiple metabolites involved in ureagenesis, including spermidine, N-acetylglutamate, glutamine, citrulline and arginine were differentially accumulated in MZ and ZZ relative to MM iHeps (**Figure 3A, Table S2**).

**Figure 3:**
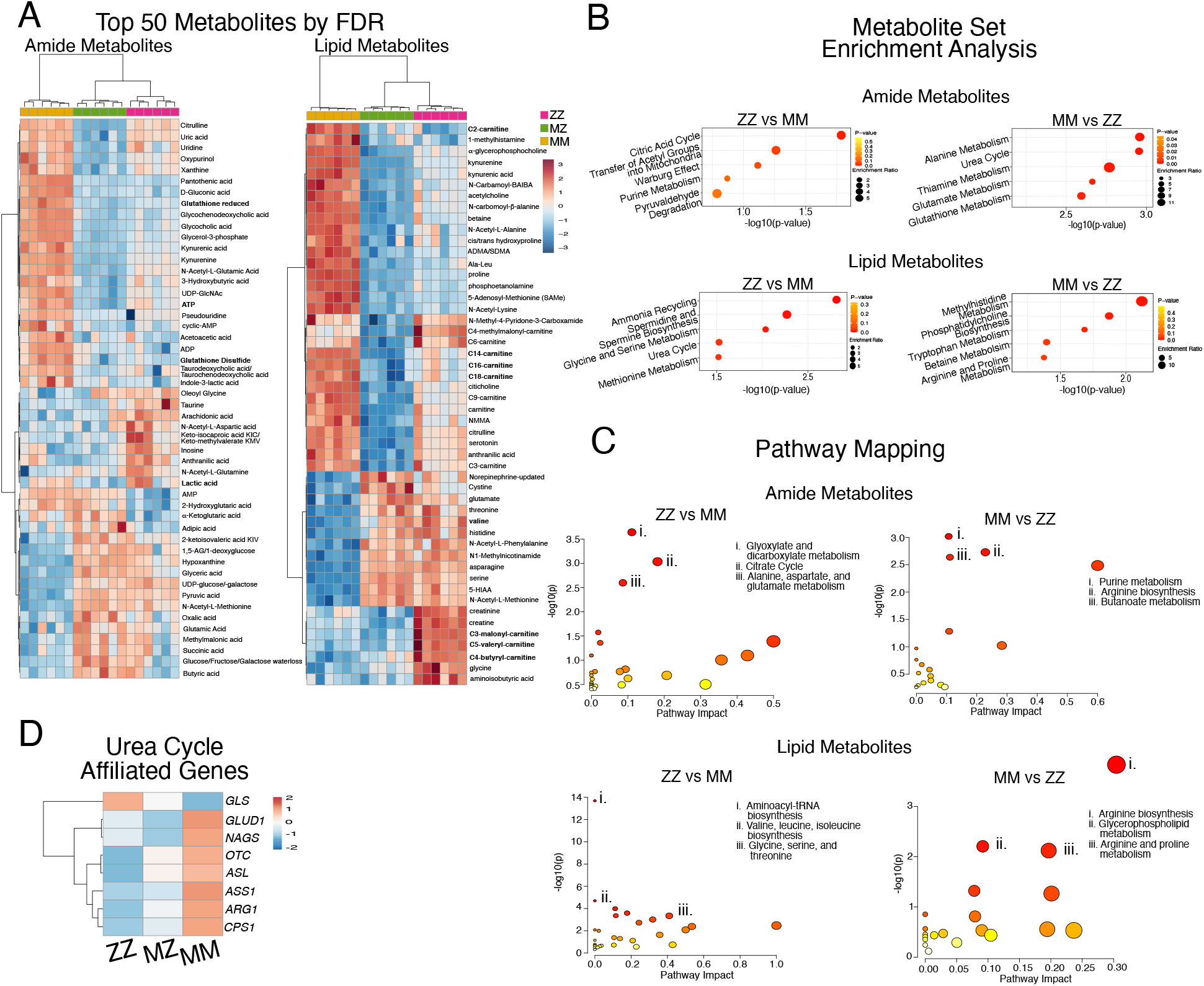
Metabolomic Analysis Identifies Dysregulated Metabolic Pathways in ZAAT-Expressing Cells. A) Heatmap of the top 50 differentially accumulated metabolites for amide and lipid fractions. B) Summary plots for metabolite set enrichment analysis for ZZ vs MM iHeps showing the top 5 pathways as ranked by p-value. C) Metabolome projection of pathway enrichment analysis for ZZ vs MM iHeps with top 3 pathways as ranked by FDR annotated. D) Heatmap of RNA expression for urea cycle enzymes.

To further evaluate metabolic pathways perturbed in ZAAT-expressing cells, we next analyzed differentially accumulated metabolites using both Metabolite Set Enrichment Analysis (MSEA) as well as Metabolic Pathway Analysis (MetPa) (33,34). We identified significant depletion (FDR < 0.05) among ZAAT-expressing iHeps in energy intensive pathways including alanine and thiamine metabolism, the urea cycle, arginine biosynthesis, which plays a key role in both the TCA and urea cycles, as well as in nicotinate and nicotinamide metabolism (NAD, NADH) (**Figures 3B,C, S2A,B**). These findings provided a functional validation of transcriptional changes in the genes encoding for urea cycle enzymes, which were decreased in ZZ or MZ iHeps compared to MM iHeps (**Figure 3D**).

The metabolite profile likewise revealed additional pathways enriched in MZ and ZZ iHeps (FDR < 0.05) including transfer of acetyl groups into mitochondria and aminoacyl-t RNA biosynthesis which, together with changes in BCAA biosynthesis and glutathione, have been associated with the induction of ER stress or the presence of chronic liver disease (**Figures 3C,D, S2A,B**)(35–37). In particular, aminoacyl-tRNA biosynthesis, the most significantly perturbed pathway among MZ and ZZ iHeps, is induced by the PERK branch of the UPR, which is activated during ER stress (36,37). Together, these data suggest that ZAAT-driven perturbations of ER and mitochondria alter homeostatic hepatic functional processes, including urea cycle and redox pathways, and potentially alter protein synthesis in both MZ and ZZ iHeps.

### ZAAT expression is associated with mitochondrial fragmentation and decreased mitochondrial respiration

To evaluate the computationally identified dysregulated mitochondrial gene expression and derangements in fatty acid oxidation in MZ and ZZ iHeps, we next imaged iHeps using transmission electron microscopy (TEM). We identified clear ultrastructural abnormalities of rough ER (rER) in both MZ and ZZ iHeps. While both MZ and ZZ iHeps exhibited dilated rER, we observed globular inclusions only within ZZ iHeps (**Figure 4A**). We noted that mitochondria in both MZ and ZZ iHeps contained distorted cristae and were significantly swollen as compared with MM iHeps (cross sectional diameter 0.416 +/- 0.011 μm, 0.339 +/- 0.004 μm, and 0.234 +/- 0.009 μm respectively) (**Figure 4A,B**). Given that swelling associated with mitochondrial respiratory defects causes mitochondria to adopt a globular shape, we calculated the mitochondrial aspect ratio (ratio of major axis:minor axis), a measure of mitochondrial fragmentation and sphericity. We found that the aspect ratio of ZZ iHeps was significantly lower than that of MM iHeps (2.202 +/- 0.20 vs 2.815 +/- 0.22 vs 3.184 +/- 0.31), while the aspect ratio of MZ iHeps fell between that of ZZ and MM iHeps but did not differ statistically (**Figure 4C**).

**Figure 4:**
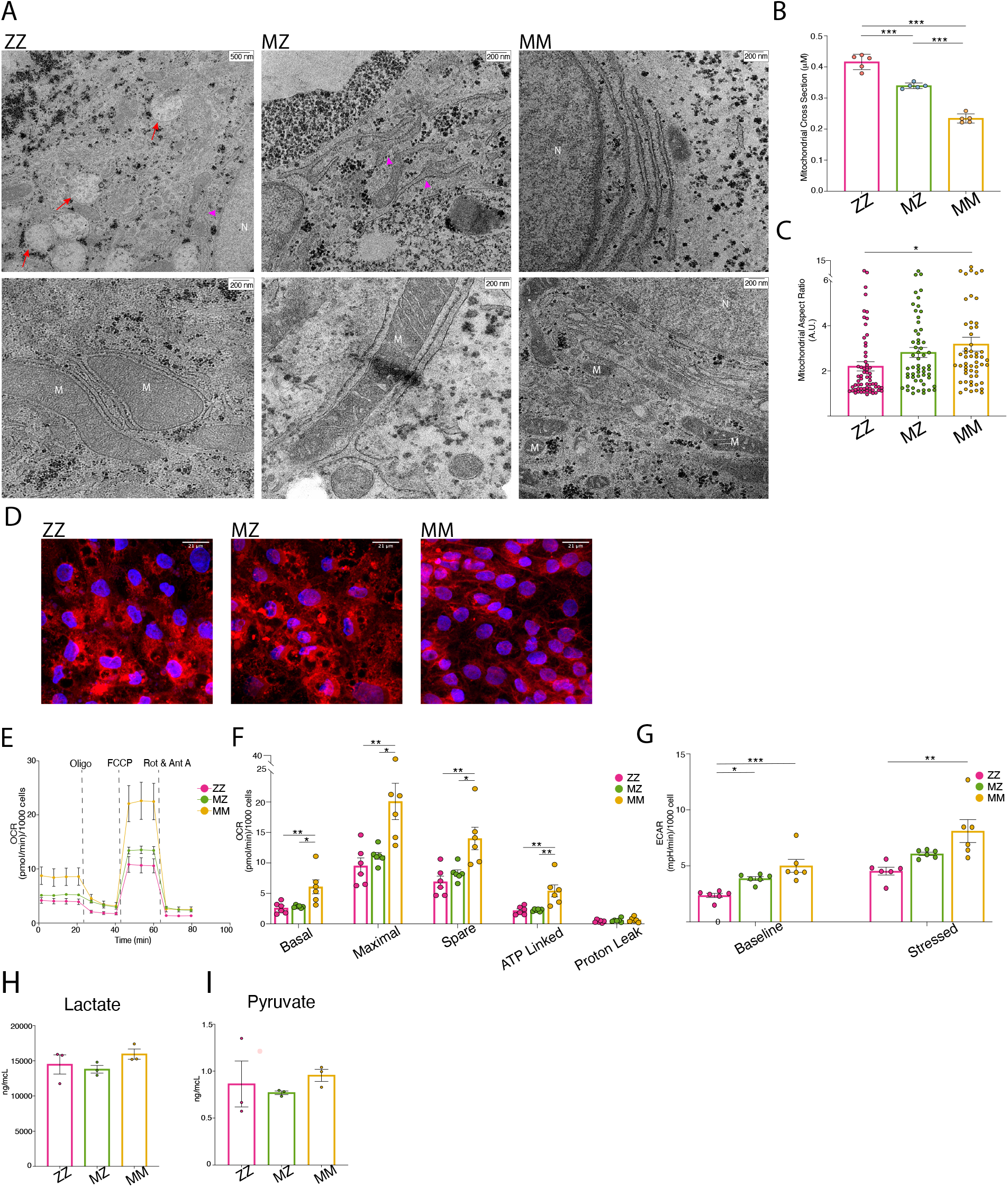
MZ and ZZ iHeps Exhibit Structural Alterations in rER and Mitochondria with Reduced Cellular Respiration. A) Transmission electron microscopy identifies rER and mitochondria in ZZ, MZ, and MM iHeps. Structural abnormalities are indicated: dilated rER (pink arrowhead), globular inclusions (red arrow). B) Average mitochondrial cross-sectional size (n=5 blinded independent observers. Mean +/- SEM). C) Average mitochondrial aspect ratio for each genotype (n=independent mitochondrial measurement. Mean +/-SEM). D) Mitotracker staining of syngeneic PiZZ6 iHeps. E) Mitochondrial oxygen consumption rate (OCR) for PiZZ6 syngeneic iHeps. F) Quantification of OCR components from (F) (n=5 independent measurements. Mean +/- SEM). G) Extracellular acidification rate (ECAR) quantification at basal and stressed states. H) Total Lactate and (I) pyruvate levels from iHep supernatants (n=3 independent experiments. Mean +/-SEM). Abbreviations: M, mitochondria; N, nucleus. *p<0.05, **p <0.01, ***P<0.001 by one-way anova with Tukey’s multiple comparison test.

To further assess mitochondrial structure, we next labeled mitochondria with MitoTracker dye. While mitochondria in MM iHeps formed an organized, tubular network (**Figures 4D, S2C**), those in MZ and ZZ iHeps were fragmented, consistent with findings noted by EM (**Figures 4D, S2C**). Having identified transcriptomic, metabolomic, and structural evidence of altered mitochondrial function in MZ and ZZ iHeps, we next performed respirometry assays to directly interrogate mitochondrial function in mutant vs corrected cells. Comparing two independent ZZ iPSC lines to their gene-corrected MZ and MM daughter lines, we observed a decrease in the basal oxygen consumption rate (OCR), as well as decreased maximal respiration capacity in MZ and ZZ relative to MM iHeps. Following the same pattern as mitochondrial fragmentation and transcriptional changes, the degree of decreased respiration was positively correlated with the number of Z mutant alleles present (**Figures 4E,F, S2D,E**). We next measured the extracellular acidification rate (ECAR) a parameter that is sensitive both to production of lactate through glycolysis as well as to oxidation of pyruvate to CO_2_. Similar to the OCR results, we found that ZZ and MZ iHeps exhibited lower ECAR at steady state, which could be explained either by reduced pyruvate oxidation to CO_2_ in the mitochondria or by decreased glycolysis production (**Figures 4G, S2F**). MM, MZ and ZZ iHeps showed similar increases in ECAR after treatment with oligomycin, which upregulates glycolysis and associated lactate excretion by blocking mitochondrial ATP synthesis and thus mitochondrial pyruvate oxidation to Acetyl-CoA. The fold increase induced by oligomycin supports decreased mitochondrial pyruvate oxidation as an explanation for reduced ECAR in MZ and ZZ cells without the need to implicate increased lactate production. To further confirm this interpretation of the OCR and ECAR data, we next quantified pyruvate and lactate in cell supernatants and found that steady state levels of these molecules did not differ based on genotype (**Figure 4H,I**). Taken together, these data demonstrate that dysmorphic mitochondria observed within MZ and ZZ iHeps exhibit aberrant function relative to genetically matched, healthy MM controls.

### scRNA-seq demonstrates transcriptional heterogeneity characterized by selective activation of UPR pathways among MZ and ZZ iHeps

While all hepatocytes express *SERPINA1*, polymerized ZAAT protein differentially accumulates within the ER of hepatocytes in ZZ patients resulting in protein “globules” that are heterogeneously distributed in vivo (3). Studies from our group and others have likewise observed variability in the amount of retained AAT among ZZ iHeps from a single preparation, based on either direct quantification by intracellular flow cytometry (**Figure 1, S1**)(19,23,24) or quantification of ER inclusion size (21). To determine whether heterogeneity similarly occurs at the transcriptional level in heterozygous ZAAT-expressing cells, we performed scRNA sequencing on syngeneic ZZ, MZ, and MM iHeps derived from derived from the PiZZ1 donor together with an additional ZZ sample from the PiZZ6 donor (PiZZ6^ZZ^) (**Figure 5**). We first compared scRNA sequencing expression data from the three PiZZ1-derived syngeneic samples to our bulk RNA sequencing results to determine whether the gene expression patterns identified by bulk RNA sequencing (**Figure 2**) were reproduced by iHeps generated from a distinct genetic donor. Applying GSEA methodology, we again found that MZ and ZZ iHeps were enriched in EMT, angiogenesis and TGF-β signaling pathways (FDR < 0.1) (**Figure S3A**).

**Figure 5:**
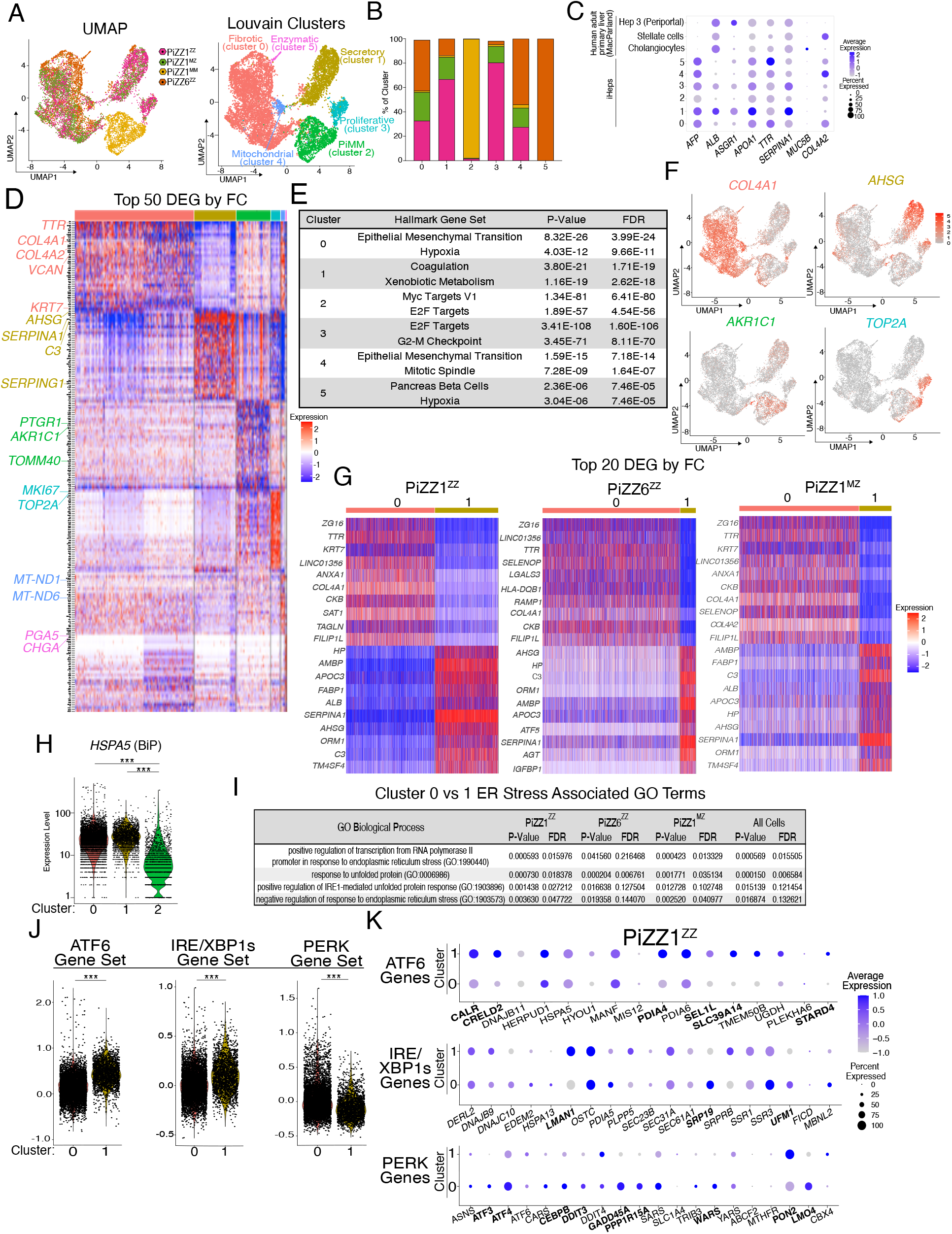
scRNA-Seq Demonstrates Transcriptional Heterogeneity Among ZAAT-Expressing iHeps with Branch-Specific Activation of the UPR. A) UMAP projection by original identity and Louvain clustering identifies 6 clusters. B) Composition of the 6 Louvain clusters by original identity. C) Average expression and frequencies of select hepatic, biliary epithelial and stellate genes across the 6 clusters. Comparison is made with a publicly available human liver single cell RNAseq dataset (62). D) Top 50 DEGs per cluster as ranked by fold change (FC) (FDR <0.001). E) Top two Hallmark gene sets for each cluster as ranked by FDR using Enrichr analysis of all DEGs (FDR<0.05). F) UMAP of select cluster specific associated DEGs. G) Top 20 DEGs as ranked by FC for the direct comparison between clusters 0 and 1 separated for each genetic background. H) Violin plot of *HSPA5* (BiP) expression by cluster. I) Top ER stress associated GO terms for cluster 0 compared to cluster 1 by original identity and for all cells. J) Violin plots for PERK, ATF6, and IRE/XBP1s module scores for clusters 0 and 1. K) Dotplot projections for UPR branch specific genes for PiZZ1^ZZ^. Differentially expressed genes are bolded (FDR <0.05). ***P<0.001

We next applied Uniform manifold approximation and projection (UMAP) as well as Louvain clustering and identified six clusters, four composed of a mixture of both ZZ and MZ cells and one composed almost entirely of MM cells (**Figure 5A,B**). Analysis of canonical hepatocyte markers demonstrated expression of a hepatic program across clusters (**Figure 5C**). Next, to identify phenotypic differences driving clustering we compared the top 50 differentially expressed genes for each cluster together with Enrichr-generated Hallmark gene set enrichments (**Figure 5D,E**) (38). Based on these analyses, we annotated the ZAAT-expressing clusters as fibrotic (cluster 0); secretory (cluster 1); proliferative (cluster 3); mitochondrial (cluster 4) and enzymatic (cluster 5). Because the MAAT-expressing cells formed a single cluster, we annotated this cluster by its genotype, “PiMM” (cluster 2). The mitochondrial and enzymatic clusters were not considered for further analysis based on their small size, and elevated expression of genes encoded within mitochondria (cluster 4) or representation of only one genetic background (cluster 5). Expression of genes associated with cell cycle and proliferation including *CKS2* and *TOP2A* distinguished both the PiMM and proliferative clusters, but the PiMM cluster additionally included genes known to be highly expressed in healthy hepatocytes, such as the aldo-keto reductase enzyme *AKR1C1* (**Figure 5D,F**). The remaining two clusters, fibrotic and secretory, contained the bulk of the MZ and ZZ cells. The fibrotic cluster contained several significantly upregulated genes associated with extracellular matrix (ECM) including *COL4A1* and *COL4A2* (**Figure 5D,F**), while the secretory cluster was characterized by expression of secreted proteins including *AHSG*, the most upregulated gene in the cluster (**Figure 5D,F**). By GSEA, the fibrotic cluster was defined by enrichment of the gene set “Epithelial Mesenchymal Transition”, while differential expression of typical hepatocyte function gene sets including “Coagulation” and “Xenobiotic Metabolism” described the secretory cluster (**Figure 5E**).

To further explore the heterogeneity observed within ZAAT-expressing iHep populations we next directly compared the fibrotic and secretory clusters. We first looked at differentially expressed genes between the two clusters, analyzing each parental line independently to avoid potential confounding resulting from genetic heterogeneity. The top differentially expressed genes defining each cluster overlapped significantly among ZZ and MZ iHeps irrespective of genetic background (**Figure 5G**) and analysis of differentially expressed genes distinguishing the two clusters demonstrated a similar pattern of Hallmark gene set enrichment with the broader cluster analysis (data not shown). ER stress and induction of the UPR are a known consequence of ZAAT expression in some but not all contexts (16,21,24,39) and defined the gene expression pattern of one cluster of ZZ iHeps in a single cell-RNA seq dataset that we recently published (23). To look for induction of the UPR among ZAAT-expressing iHeps in this study, we first evaluated expression of the ER chaperone *HSPA5* (BiP) to see if its expression varied among clusters. While BiP expression was significantly higher in ZAAT-expressing clusters relative to the PiMM cluster (cluster 2) (FDR <0.001), it did not differ significantly between the ZAAT fibrotic (cluster 0) and secretory (cluster 1) clusters (**Figure 5H**). We next performed Gene Ontology enrichment analysis and identified enrichment for terms related to ER stress and the UPR relative among MZ and ZZ iHeps within the fibrotic cluster compared to those in the secretory cluster (**Figure 5I**). To further explore whether UPR activation might differ between clusters, we applied gene modules specific for each of the three branches of the UPR (ATF6, PERK, IRE1/XBP1s) (40). Intriguingly, gene sets defining specific UPR branches were, in fact, differentially expressed between the two clusters (**Figures 5J,K, S3B**). While cells in the secretory cluster exhibited significantly higher expression of genes associated with ATF6 and IRE1/XBP1s activation, cells in the fibrotic cluster expressed higher levels of genes associated with activation of PERK (**Figures 5J,K, S3B**). To further validate these findings, we applied the same analysis to an additional scRNA seq dataset including ZZ and MM iHeps derived from a 3^rd^ distinct iPSC line, PiZZ100 (**Figure S3C**). Despite using different sequencing platforms in the two experiments, we observed differential expression of UPR branch-specific genes among two ZZ clusters. These two ZZ clusters were defined by enrichment of Hallmark gene sets overlapping with the fibrotic and secretory clusters, as well as being distinguished by the expression of genes associated with either PERK (fibrotic) or ATF6 and IRE1/XBP1s (secretory) activation (**Figure S3C-G**). Together, these data demonstrate the presence of transcriptional heterogeneity within mutant ZAAT-expressing iHeps characterized by the expression of profibrotic genes in a subset of MZ and ZZ cells. This aberrant expression pattern is further accompanied by a relative downregulation of the ATF6 and IRE1/XBP1s UPR branches, together with upregulation of PERK effector genes.

## Discussion

Our studies demonstrate that expression of a single Z allele is sufficient to significantly perturb intracellular AAT protein processing in iPSC derived-hepatocytes and to induce morphologic and functional derangements of both ER and mitochondria with associated transcriptomic and metabolomic changes that overlap significantly with those observed in homozygous mutant ZZ cells.

Aggregated polymers of misfolded ZAAT protein that accumulate in and distort the ER, are a hallmark of the disease and have been identified previously in ZAAT-expressing cell lines, PiZ mice, ZZ iHeps, and ZZ and MZ human tissue specimens (5,16,21,22,24,41). Damaged mitochondria with morphological distortion and defective mitophagy have similarly been observed in ZZ patient iHeps and liver tissue (22,41). Our studies extend those previous observations to heterozygous MZ cells, in which we observed gross structural alterations of mitochondria and ER that increased in severity with increasing Z allele copy number and were associated with diminished basal and maximal cellular respiration. We further identified evidence of mitochondrial fragmentation in ZZ but not MZ iHeps, potentially consistent with a correlation between the severity of ER stress and protein misfolding and the degree of mitochondrial injury.

In addition to structural differences, we identified alterations of the global transcriptome and metabolome in both MZ and ZZ iHeps, some of which have been previously associated with ER stress or mitochondrial dysfunction (35–37,42). Among these were enrichment for metabolites significant in the citric acid cycle, glutathione metabolism, and branched chain amino acid and tRNA biosynthesis. Analysis of both transcriptomic and metabolomic data suggested that multiple metabolic pathways were dysregulated in iHeps that carry even a single copy of the mutant Z allele. Urea cycle metabolites were significantly altered in both MZ and ZZ iHeps with associated downregulation of urea cycle enzymes (ASS1, CPS1, OTC, ASL) and their transcriptional regulators, HNF4a, HNF1a, and CEBPA, supporting previous observations in PiZ mice of ureagenesis impairment (43). Indeed, HNF4α has also been shown to stimulate mitochondrial biogenesis and its downregulation explained mitochondrial dysfunction in other forms of liver disease (44). While the mechanisms underlying these findings have not been delineated, Piccolo et al speculate that downregulation of HNF4a results as a downstream consequence of NF-κB activation in response to accumulated ZAAT. Further studies will be needed to determine whether this hypothesis explains experimental findings in iHeps.

It has long been observed that aggregated ZAAT “globules” are heterogeneously distributed in patient liver biopsy specimens (3). These findings have been echoed in ZZ patient iHep studies noting heterogeneity of total cellular ZAAT content and ER inclusion size (19,21,23,24). To better understand the transcriptional basis for these findings, we analyzed approximately 12,000 ZZ, MZ, and MM iHeps by scRNA sequencing. Intriguingly, while MM cells formed a single heterogenous cluster driven predominantly by genotype, ZAAT-expressing iHeps formed two transcriptionally distinct clusters one of which was characterized by expression of pro-fibrotic and ER stress-associated genes with the other distinguished by expression of proteins classically secreted by hepatocytes (*AHSG*, *C3 and SERPING1*). Further analysis then revealed differential activation of specific branches of the UPR between the two clusters, with upregulation of the ATF6 and IRE1/XBP1s arms in the secretory cluster and downregulation of these arms together with increased expression of PERK branch components in the pro-fibrotic/ER stress cluster. This heterogeneity further informs understanding of the hepatic response to misfolded protein aggregates in AATD and offers a potential explanation for previous observations that have not identified UPR activation via bulk sequencing methodologies (16,21,24,39). These data are likewise consistent with prior literature demonstrating that heterogenous activation of the UPR can exist in an apparently homogenous cell population (45). Activation of PERK with associated expression of cell stress/apoptotic mediator CHOP (*DDIT3*) has been shown to occur in the context of prolonged UPR activation (46,47). Whether the relative inactivation of the ATF6 and IRE1 branches and activation of PERK genes among cellular subsets results from a kinetic of gene expression or is associated with an increased burden of intracellular ZAAT remains unclear. Further studies, potentially including single cell analysis of human liver tissue, will be useful to provide additional context to these observations.

This study utilizes genetically edited patient iPSCs to control for genetic heterogeneity and identify cellular phenotypes that result from expression of either one or two mutant ZAAT alleles. While population-based studies have identified increased liver disease risk among MZ individuals, it remains likely such risk is heterogenous in the population and possibly modulated by unidentified genetic factors affecting either cellular degradation pathways or cellular response to accumulated protein aggregates as has previously been hypothesized (48–50). Further studies will be needed to further explore the mechanisms by which intracellular accumulation of ZAAT leads to the metabolic dysregulation observed here, to delineate their generalizability among cells derived from individuals with distinct disease outcomes, and to determine whether they can ultimately be interrogated to quantify potential risk in MZ or ZZ individuals before they develop disease.

In summary, our studies demonstrate that MZ iHeps exhibit a cellular phenotype intermediate to genetically matched ZZ and MM comparators. Through multi-omic profiling, we have shown that AAT processing is deranged in both ZZ and MZ iHeps and associated with downstream metabolic dysregulation, impaired mitochondrial function, and cellular heterogeneity characterized by branch-specific UPR gene expression. These findings provide important insight into the mechanistic underpinnings of ZAAT-driven hepatocyte injury that could contribute to the increased risk of clinical liver disease observed among MZ and ZZ individuals.

## Methods

### iPSC Line Generation and Maintenance

All experiments involving the differentiation of human iPSC lines were performed with the approval of Boston University Institutional Review Board (BUMC IRB protocol H33122). The three parental PiZZ iPSC lines (PiZZ1, PiZZ6 and PiZZ100) have been previously published (19,24). iPSCs were maintained in feeder-free conditions on growth factor reduced Matrigel (Corning) in mTeSR-1 media (StemCell Technologies) using either gentle cell dissociation reagent (GCDR) (StemCell Technologies) or ReLeSR (StemCell Technologies). All iPSC parental and gene-edited lines were verified to be free of mycoplasma and karyotypically normal as determined by G-band karyotyping analysis from 20 metaphases. Further details of iPSC derivation, characterization and culture are available for free download at https://www.bu.edu/dbin/stemcells/protocols.php. iPSCs utilized in this manuscript are available upon request from the CReM iPSC repository at https://stemcellbank.bu.edu.

### CRISPR Based Editing of Z Mutation

The CRISPR/Cas9 endonuclease system was used to target the *SERPINA1* sequence in close proximity to the Z mutation site within exon 5 using a previously published protocol (19,51). To achieve scarless editing and generation of both mono- and bi-allelic corrected iPSCs, two 70bp ssODN repair templates were synthesized (Integrated DNA Technologies) with sequence homology to the *SERPINA1* locus adjacent to the Z mutation and in the complimentary orientation with respect to the gRNA sequence (25). The donor template includes either the wild type *SERPINA1* sequence (T to **C**) or retained the Z mutation (**T**) sequence. Both ssODN’s contained a silent mutation (G to **A**) to insert a new ClaI restriction enzyme digest site and facilitate screening for template incorporation. Two additional silent mutations (C to **T** and C to **A**) were included for the M-donor, with the C to T mutation introduced to reduce subsequent retargeting by Cas9. For the Z-Donor the C to A silent mutation was not included. The resulting M-ssODN sequence was: 5’ TCT AAA AAC ATG GCC CCA GCA GCT TCA GT**A** CCT TT**T** T**CA** TCG ATG GTC AGC ACA GCC TTA TGC ACG GCC T 3’; and the Z-ssODN sequence: 5’ TCT AAA AAC ATG GCC CCA GCA GCT TCA GTC CCT TT**T** T**TA** TCG ATG GTC AGC ACA GCC TTA TGC ACG GCC T 3’ (52). When iPSCs were in log phase growth they were pretreated with 10μM Y-27632 (Tocris) for 3h. Cells were dissociated into single cell suspension with GCDR, and 5×10^6^ cells were resuspended in 100μL P3 solution containing Supplement1 (Lonza) with 5μg plasmid DNA (containing gRNA and Cas9-2A-GFP), and 5 μg of each ssODN donor (10μg total). The cell/DNA mixture was then nucleofected using the 4-D nucleofector system (Lonza) code CB-150 and densely replated on Matrigel coated 6-well plates at 2.5×10^6^ cells per well. After 48h, cells were again pretreated with 10μm Y-27632 for 3h before GCDR was used to achieve single cell suspension and GFP+ cells were isolated using a MoFlo Legacy cell sorter (Beckman Coulter). Sorted GFP+ single cells were sparsely replated at 1×10^4^ cells/well in a 10cm Matrigel coated dish to facilitate clonal outgrowth. Immediately post-sort, cells were grown in recovery media consisting of 1-part mTeSR-1 and 1-part iPSC conditioned mTeSR-1 supplemented with 0.7ng/mL FGF2 until multicellular colonies could be observed at which time they were grown in mTeSR-1 (53). For the first 24h, 10μm Y-27632 was added to recovery media. After approximately 10 days, emergent colonies were of sufficient size for individual selection, expansion, and screening using the following PCR primers: 5’ GCA GAC GTG GAG TGA CGA TG 3’ and 5’ CCT GGA TTC AAG CCC AGC AT 3’. PCR product was then assessed using a ClaI restriction enzyme digest (New England BioLabs) per manufacturer’s protocol. Positive digest resulted in 2 bands of 298bp and 407bp. Digested clones were further characterized by Sanger sequencing to confirm donor template incorporation.

### iHep Generation

iPSC directed differentiation to the hepatic lineage was performed using our previously published protocol (19,24). Briefly, undifferentiated iPSCs were cultured until confluent then passaged at Day 0 using GCDR, replated at 1×10^6^ cells per well of a Matrigel coated 6 well plate, and placed into hypoxic conditions (5% O2, 5%CO_2_, 90%N_2_) for the remainder of the differentiation. They were patterned into definitive endoderm using the STEMdiff Definitive Endoderm Kit per manufacturer’s instructions over 5 days (StemCell Technologies). Cells were passaged on day 5 of differentiation using GCDR, with endoderm efficiency confirmed via cell surface staining for CXCR4 and cKit, and endoderm subsequently grown in serum free base media supplemented with stage specific growth factors to specify the hepatic lineage and induce maturation (**Figure 1B**). Detailed protocols for derivation of iPSC-hepatocytes are available for free download at: https://crem.bu.edu/cores-protocols/.

### Flow Cytometry

Endoderm induction was quantified using anti-human CD184(CXCR4)-PE (StemCell Technologies) and anti-human CD117(CKIT)-APC (ThermoFisher Scientific) conjugated monoclonal antibodies (19,24). To quantify intracellular protein content, iHeps were fixed in 1.6% paraformaldehyde for 20 min at 37°C and then permeabilized in saponin buffer (BioLegend). Cells were first probed using AAT (Santa Cruz Biotechnologies), AFP (Abcam), ZAAT (a kind gift from Qiushi Tang and Chris Mueller), and then anti-mouse IgG1-AlexaFluor647 (Jackson ImmunoResearch), anti-rabbit IgG-AlexaFluor488 (ThermoFisher Scientific) and anti-mouse IgG-AlexaFluor488 (ThermoFisher Scientific) antibodies. Staining quantification was performed using BD FACSCalibur or Stratedigm S1000EXi and all gating was performed using isotype-stained controls. Data analysis was performed using FlowJo (Tree Star) and Prism8 (GraphPad) software.

### ELISA

Secreted total AAT was quantified from iHep supernatants using the human alpha-1-antitrypsin ELISA quantification kit (GenWay Biotech) per manufacturer’s instructions. ZAAT ELISA was performed by adapting this protocol to incorporate the anti-ZAAT antibody and ZAAT standard (23,27).

### Neutrophil Elastase Inhibition

To quantify the capacity of iHep secreted AAT to inhibit neutrophil elastase, 4ml of iHep supernatant was concentrated using Amicon Ultra-4 MWCO 30kDa spin columns (MilliporeSigma). Serial dilutions of the concentrated supernatants were then incubated with human neutrophil elastase (MilliporeSigma) in the presence of methoxysuccinyl-Ala-Ala-Pro-Val-p-nitroanilide (Millipore Sigma). To quantify bioactivity colorimetric change was quantified using a 96 well plate reader set at 405nm (54).

### Immunohistochemistry

For immunohistochemistry analysis, cells were passaged at day 5 of differentiation into 2-well chamber slides for the remainder of hepatic directed differentiation (ThermoFisher Scientific). For mitochondrial morphology analysis MitoTracker Deep Red FM (ThermoFisher Scientific) was applied at 200nM for 45 min prior to fixation. Cells were fixed using 4% paraformaldehyde for 10 min at room temperatures. For antibody labeling, cells were permeabilized with 0.3% Triton-X (MilliporeSigma) and blocked with 4% normal donkey or goat serum prior to incubation overnight with primary antibodies. Cells were probed using anti-AAT (Santa Cruz Biotechnologies), HNF4A (Abcam), or 2C1 (a kind gift from Elena Miranda and David Lomas) antibodies. Following incubation with primary antibody, cells were washed in PBS with 0.05% Tween20 (MilliporeSigma) and then incubated with secondary antibodies, anti-rabbit IgG-AlexaFluor647 and anti-mouse IgG-AlexaFluor488, for 1h at room temperature. Finally, cells were again washed with PBS containing 0.05% Tween20 and then counterstained with Hoechst 3342 (ThermoFisher Scientific). Cells were imaged using the Zeiss LSM 710-Live Duo scan confocal microscope and images were processed with ImageJ software.

### AAT Pulse Chase Radiolabeling

AAT secretion kinetics were assayed via pulse-chase radiolabeling. The day prior to labeling fully differentiated iHeps were seeded onto 24 well plates to achieve at least 90% confluency. Radiolabeling was then performed using previously published methods (22,55). Briefly, cells were incubated for 1h in methionine (Met)- and cysteine (Cys)-free DMEM supplemented with 250μCi of ^35^S-Met/Cys radiolabel (PerkinElmer) per condition at normoxia. Cell were then washed and refed with Met/Cys containing DMEM for the 4h chase period. At time 0 and every hour supernatants and cell lysates were harvested. AAT was then immunoprecipitated from lysates and supernatants using a polyclonal antihuman AAT antibody (Proteintech) and resolved by 10%v/v Tris-Glycine polyacrylamide gel fluorography (ThermoFisher Scientific). Densitometric analysis was performed using ImageJ with the relative densitometric value of T0 set at 100%. (protocols generously provided by Ira Fox, University of Pittsburgh and David Perlmutter, Washington University School of Medicine in St. Louis).

### RNA Sequencing

Total RNA from PiZZ6 syngeneic ZZ, MZ and MM iHep differentiations were isolated in triplicate using the miRNeasy kit (Qiagen) per manufacturer’s instruction. mRNA was then isolated using magnetic bead-based poly(A) selection followed by synthesis of cDNA fragments. cDNA fragments were then end-paired, and PCR amplified to create each cDNA library. Sequencing was performed using the NextSeq 500 (Illumina) with a post sequencing Phred quality score >90%. Reads were then aligned to the ENSEMBL human reference genome GRCh38.91 using *STAR* (56,57). We used the Bioconductor package *edgeR* to import, filter and normalize the count matrix, followed by the *limma* package and *voom*, for linear model fitting and differential expression testing, using empirical Bayes moderation for estimating gene-wise variability prior to significance testing based on the moderated t-statistic (58–60). We used a FDR corrected p-value of 0.05 as threshold to call differentially expressed genes (61). Gene set analysis was later performed by querying the enrichment of gene sets from the Hallmark collection and separately any gene set from the GO collection that was related to mitochondria, ER and ER stress (31). The same analysis was subsequently applied to an external data set generated by an independent laboratory and openly available on ArrayExpress (accession number E-MTAB-6781).

### scRNA Sequencing

For single cell RNA sequencing (scRNA-seq) PiZZ1 syngeneic (ZZ, MZ, MM) and PiZZ6 ZZ iHeps were disassociated using 0.25% Trypsin at day 25 of differentiation and sorted for live cells using Calcein Blue (ThermoFisher Scientific) on a MoFlo Astrios EQ (Beckman Coulter). Single live cells were then captured, and library preparation performed using the Chromium Single Cell 3’ v3 user protocol and Chromium Controller instrument per manufacturer instructions (10X Genomics). Each library was then sequenced using the Illumina NextSeq 500 to obtain sequencing depths of between 25-50K reads/cell. Fastq and count matrix files were generated using Cell Ranger v 3.0.2 and the transcriptome mapped again to the ENSEMBL human reference genome GRCh38. We then used Seurat v3 to further process and analyze the data. Data was normalized using the regularized negative binomial regression method with cell degradation regressed out. We then performed dimensionality reduction using the uniform manifold approximation and projection (UMAP) to represent the data and for clustering the Louvain algorithm was utilized. Differential gene expression was determined by a log fold change of 0.25 using the Wilcoxon rank-sum test, and GSEA was performed using hypeR. UPR arm specific module score significance was determined using Welch Two Sample t-tests. For comparison of canonical hepatocyte genes to primary human hepatocytes the dataset was merged with a publicly available scRNA-seq dataset generated by an independent lab available at NCBI GEO accession GSE115469 (62).

A second independent scRNA-seq analysis was performed using PiZZ100 ZZ and MM iHeps as described above except a large (15-25μm cell diameter) 800 cell capacity microfluidic chip for mRNA-seq (Fluidigm) was utilized for single cell capture. After a live-dead sort cells were loaded onto the chip per manufactures instructions using the Fluidigm C1 HT workflow to capture, lyse, reverse transcribe RNA and for library preparation.

### Metabolomics

Day 26 PiZZ6 ZZ, MZ and MM iHep media was aspirated from 6 well plates and cells quickly washed using high performance liquid chromatography (HPLC) grade water. The culture plate was then carefully inverted in a liquid nitrogen-resistant basin and liquid nitrogen was poured directly onto the plate bottom to quench cellular metabolic activity. Liquid nitrogen was allowed to evaporate (30-60s) before 2mLs of extraction medium consisting of 75% (9:1 v/v) Methanol (ThermoFisher Scientific):Chloroform (ThermoFisher Scientific) mixture, 25% water were poured into each well. Cells were then lifted from the plate and 1mL was transferred to an Eppendorf tube, and 1mL transferred into a collective pooled extract tube. Extracts were centrifuged at 16,000 rpm for 5 min and split for Amide and Lipid analyses. Labeled isotope standards L-Phenylalanine-d8 (Sigma-Aldrich) and L-Valine-d8 (Cambridge Isotope Laboratories) were added to the supernatants. Samples were then dried down on a speedvac concentrator (ThermoFisher Scientific) and re-suspended in 100 mL of (50:50 v/v) acetonitrile (J.T. Baker): water before injection. Sample injection volume was 5 or 10 mL, depending on liquid chromatography-tandem mass spectrometry (LC-MS) acquisition method, described below.

LC-MS data were acquired using two methods on two LC-MS machines. The Lipid method, was acquired using a 4000 QTrap triple quadrupole mass spectrometer (Applied Biosystems/Sciex) that was coupled to a multiplexed LC system comprised of a 1200 Series pump (Agilent Technologies) and an HTS PAL autosampler (Leap Technologies) equipped with two injection ports and a column selection valve. Cellular lipid extracts were analyzed using a 150 mm x 3.2 mm Phosphere C4 column (Grace) and mobile phases (mobile phase A: [95:5:0.1 v/v/v] 10mM Ammonium Acetate (Sigma-Aldrich): Methanol: Acetic Acid (Sigma Aldrich), mobile phase B: 0.1% Acetic acid (Sigma-Aldrich) in Methanol). A 10 mL volume of extract was injected directly onto the column under initial conditions (80:20 Mobile Phase A: Mobile Phase B, with a 350 mL/min flow rate). The solvent composition was held constant for 2 min, followed by a linear gradient to 20:80 A:B over 1 min, and a further linear gradient to 0:100 A:B over 12 min, where it was held for 10 min before returning to initial conditions for a 10 min re-equilibration. The LC system was connected to an API-4000 QTrap triple quadrupole mass spectrometer (Applied Biosystems) run in positive ion mode. MS ionization was achieved using an electrospray ionization (ESI) source. Ions were measured in full scan mode (Mass range: 400-1100 Daltons, Dwell time: 1.25 s / scan for a total of 1099 scans / sample). Metabolite peaks were integrated using Multiquant Software (AB Sciex).

The Amide method used HILIC chromatography on a 2.1 x 100mm 3.5 μm Xbridge Amide column (Waters) in negative ion mode. Mobile phase A was (95:5 v/v) water: acetonitrile with 20 mM ammonium acetate (Sigma-Aldrich) and 20mM ammonium hydroxide (Sigma-Aldrich) (pH 9.5). Mobile phase B was acetonitrile. The chromatography system consisted of a 1260 Infinity autosampler (Agilent Technologies) connected to a 1290 Infinity HLPC binary pump system (Agilent Technologies). Injection volume was 5 μL. The initial conditions were 0.25 mL/min of 85% mobile phase B followed by a linear gradient to 35% mobile phase B over 6 min. This was followed by a linear gradient to 2% mobile phase B over 30s held for an additional 30s, then a 30s gradient return to 85% mobile phase B. Column equilibration was continued for 4.5 min at 0.5 mL/min for a total cycle time of 12 min. The column compartment was maintained at 30 Celsius. The HPLC pump was connected to a 6490 QQQ (Agilent Technologies) triple quadrupole mass spectrometer equipped with an electrospray ionization source, on which 157 metabolites were optimized for negative mode detection. Final mass spectrometry settings for the QQQ 6490 were sheath gas temperature 400 Celsius, sheath gas flow 12 L/min, drying gas temperature 290 Celsius, drying gas flow 15 L/min, capillary voltage 4000V, nozzle pressure 30 psi, nozzle voltage 500V, and delta EMV 200V. Metabolite quantification was determined by integrating peak areas using MassHunter QQQ Quant (Agilent Technologies).

For both methods, all metabolite peaks were manually reviewed for peak quality. In addition, pooled cellular extracts were run every 10 injections, enabling the monitoring and correction for temporal drift in mass spectrometry performance. All samples were normalized to the nearest pooled sample in a metabolite-by-metabolite manner. Metabolites were then uploaded and normalized using MetaboAnalyst v5 (33,34). Differential metabolites were identified using an ANOVA at an FDR <0.05 and for pairwise differences a post-hoc Fisher’s LSD was used. Metabolite Set Enrichment Analysis (MSEA) as well as Metabolic Pathway Analysis (MetPa) were then performed on differential metabolites with significant enrichment determined at an FDR < 0.05.

### Transmission Electron Microscopy

PiZZ6 ZZ, MZ, and MM iHeps were prepared using a published publicly available protocol (22). iHeps were fixed for 1h in 2.5% glutaraldehyde (Ladd Research) in PBS pH 7.4 at room temperature. The samples were then washed three times in PBS. Post fixation samples were then placed in 1% osmium tetroxide and 1% potassium ferricyanide in PBS for 1h at room temperature in the dark. Samples were then dehydrated in a step wise fashion using 30%, 50%, 70& and 90% ethanol for 10 min followed by 100% ethanol for 15 min three times. Next, the sample was changed to EMbed 812 (EMS) for 1h three times. Finally, samples were embedded in fresh Embed 812 by placing an inverted BEEM capsule onto cultures and polymerized overnight at 37 Celsius then 48h at 60 Celsius. Plastic embedded samples were then thin sectioned at 70nm and grids stained in 4% aqueous Uranyl Acetate for 5min at 60 Celsius followed by lead citrate for 10 min at room temperature. Sections on grids were imaged using a Philips CM12 EM operated at 100kV, and images were recorded on a VIPS F216 CMOS camera with a pixel size of 0.85-3.80 nm per pixel.

### Lactate and Pyruvate Quantification

To quantify extracellular levels of lactate and pyruvate from iHep supernatants aliquots were taken at day 22 of differentiation and analyzed using Lactate and Pyruvate Assay Kits (Sigma-Aldrich) per manufacturer’s instructions. Colormetric change was quantified using the Infinite M1000 Pro Plate Reader (TECAN).

### Respirometry Assays

Day 20 ZZ, MZ and MM iHeps were seeded onto 96 well Matrigel coated Seahorse XF Cell Culture Microplate (Agilent Technologies) at a density of 50,000 cells/well. After 24h fresh media was applied for another 24h. The Agilent Seahorse XF Cell Mito Stress Test was then performed per manufacturer’s instructions. Briefly, iHeps were washed two times with prewarmed Seahorse Media (XF Base with 25mM Glucose and 10mM Pyruvate added), 180μL of assay media added, and cells were placed into a non-CO2 37 Celsius incubator for 60 min. The prepared sensor cartridge and cells were then loaded into the Agilent Seahorse XFe96 Extracellular Flux Analyzer for Oxygen Consumption Rate (OCR) and Extracellular Acidification Rate (ECAR) quantification. Port injections contained the following: Port A, oligomycin at a final concentration of 2μM; Port B, FCCP at a final concentration of 1μM; Port C, antimycin A and rotenone at a final concentration of 1μM. To normalize for cell number iHeps were fixed in paraformaldehyde for 10 min at room temperature then stained with Hoechst 3342. Widefield images of each well were then obtained using the Nikon IHC microscope and individual nuclei quantified using ImageJ.

### Statistics

Statistical methods for each assay are indicated in appropriate figure legend and individual methods sections and differences were considered significant at an adjusted p < 0.05. Bioinformatic data was adjusted for multiple hypothesis testing using Benjamini-Hockberg procedure and FDR <0.05 was used to indicate significance unless otherwise indicated.

## Study Approval

The IRB of Boston University approved the generation of all iPSC samples with documented informed consent obtained prior to obtaining any human samples.

## Authors Contributions

J.E.K. and A.A.W. contributed to conception and design of the study. J.E.K., R.B.W., T.M., M.I.H., M.D., A.H., E.B., X.S., R.E.G., A.N.H., and A.A.W. contributed to data collection. J.E.K., R.B.W., F.W., T.M., J.L.-V., E.B., I.S.C., N.B.-P., M.L., A.N.H., D.N.K, and A.A.W. contributed to data analysis. J.E.K., D.N.K., and A.A.W. contributed to manuscript preparation.

## Acknowledgments

The authors would like to thank members of the Wilson and Kotton labs at the Center for Regenerative Medicine (CReM) for helpful scientific discussions. We thank Greg Miller, CReM Laboratory Manager, Marianne James, CReM iPSC Core Manager, and Meenakshi Lakshminarayanan, CReM Administrative Director, for their invaluable support. Special thanks to David Perlmutter and members of his laboratory, Tunda Hidvegi and Pam Hale, for training in pulse-chase radiolabeling. For technical support we would like to thank Brian R. Tilton of the Boston University Flow Cytometry Core Facility, Yuriy Aleksyeyev of the Sequencing Core Facility, Lynn Deng and Matthew Au of the Analytical instrument Core, and Michael T. Kirber of the Cellular Imaging Core. Figure schematics and cartoons were created with BioRender.com. Finally, we thank the Alpha-1 community as without their selfless participation none of this work could be done. This work was supported by an Alpha-1 Foundation John W. Walsh Translational Research Award (to J.E.K.), CJ Martin Early Career Fellowship from the Australian National Health and Medical Research Council (to R.B.W), NIH grant R01HL095993 (to DNK), iPSC distribution and disease modeling is supported by NIH grants U01TR001810 (to D.N.K and A.A.W) N0175N92020C00005 (to D.N.K.), The Alpha-1 Project (TAP)-a wholly owned subsidiary of the Alpha-1 Foundation (to D.N.K and A.A.W) and NIH grants R01DK101501 (to A.A.W.), and R01DK117940 (to A.N.H. and A.A.W).

