## Supplemental Figures for "Modeling of Alpha-1 Antitrypsin Deficiency with Syngeneic Human iPSC-Hepatocytes Reveals Metabolic Dysregulation and Cellular Heterogeneity in PiMZ and PiZZ Hepatocytes"

### Supplemental Material

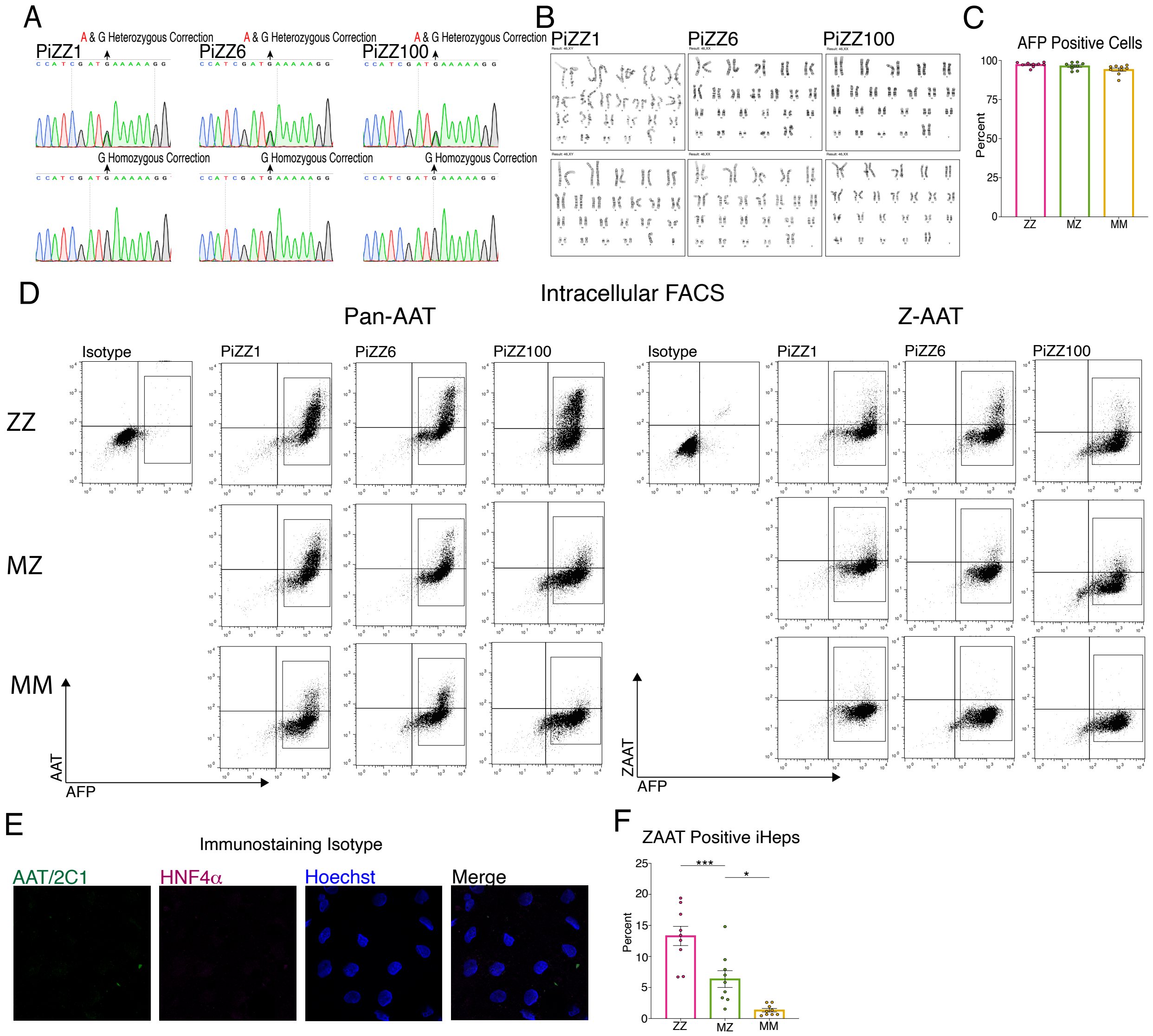


**Figure S1: Characterization of MZ and MM CRISPR/Cas9 generated iPSCs and iPSC derived hepatic cells**. A) Sanger sequencing for each of the MZ and MM iPSCs. B) Each line was confirmed to have retained a normal karyotype following CRISPR targeting. C) Percent of iHeps staining positive for AFP (Mean +/- SEM) D) Representative FACS plots for PiZZ1, PiZZ6, or PiZZ100 ZZ, MZ, or MM iHeps stained with pan-AAT or ZAAT antibodies together with antibodies against AFP. E) Immunostaining isotype controls for AAT, 2C1 and HNF4α antibodies. F) Percent ZAAT positive iHeps identified in C (mean± SEM). *p<0.05, ***p<0.001 by one-way anova with Dunnett's multiple comparisons.

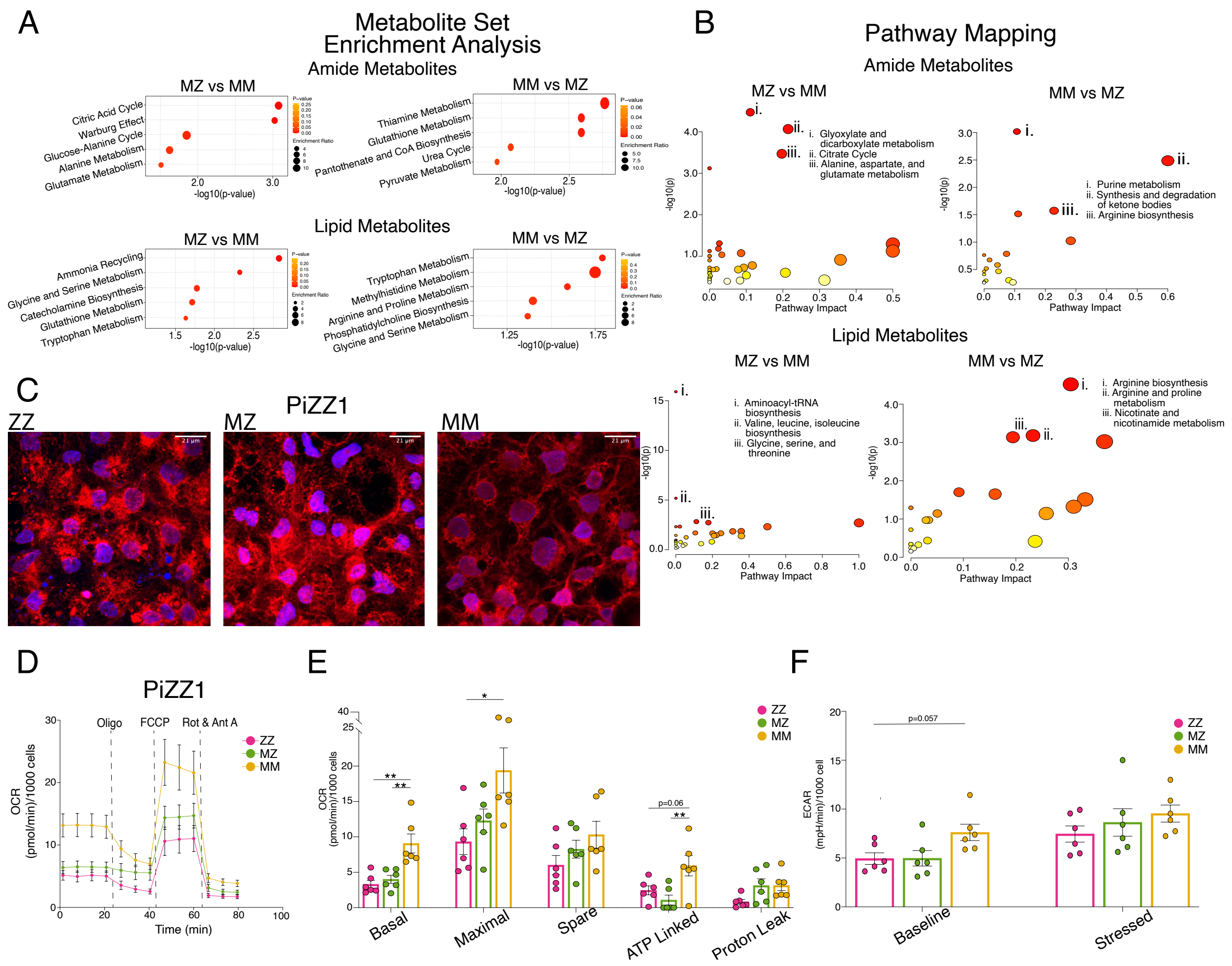


**Figure S2: MZ and ZZ iHeps Demonstrate Metabolic and Mitochondrial Dysregulation**. A) Summary plots for metabolite set enrichment analysis from MZ vs MM iHeps showing the top 5 pathways as ranked by p-value. B) Metabolome projection of pathway enrichment analysis for MZ vs MM iHeps with top 3 pathways as ranked by FDR annotated. C) Mitotracker staining of PiZZ1 syngeneic iHeps. D) Mitochondrial oxygen consumption rate (OCR) for PiZZ1 syngeneic iHeps. E) Quantification of OCR components from (D). G) Extracellular acidification rate (ECAR) quantification at basal and stressed states. *p <0.05, **p<0.01 by one-way anova with Tukey's multiple comparison test.

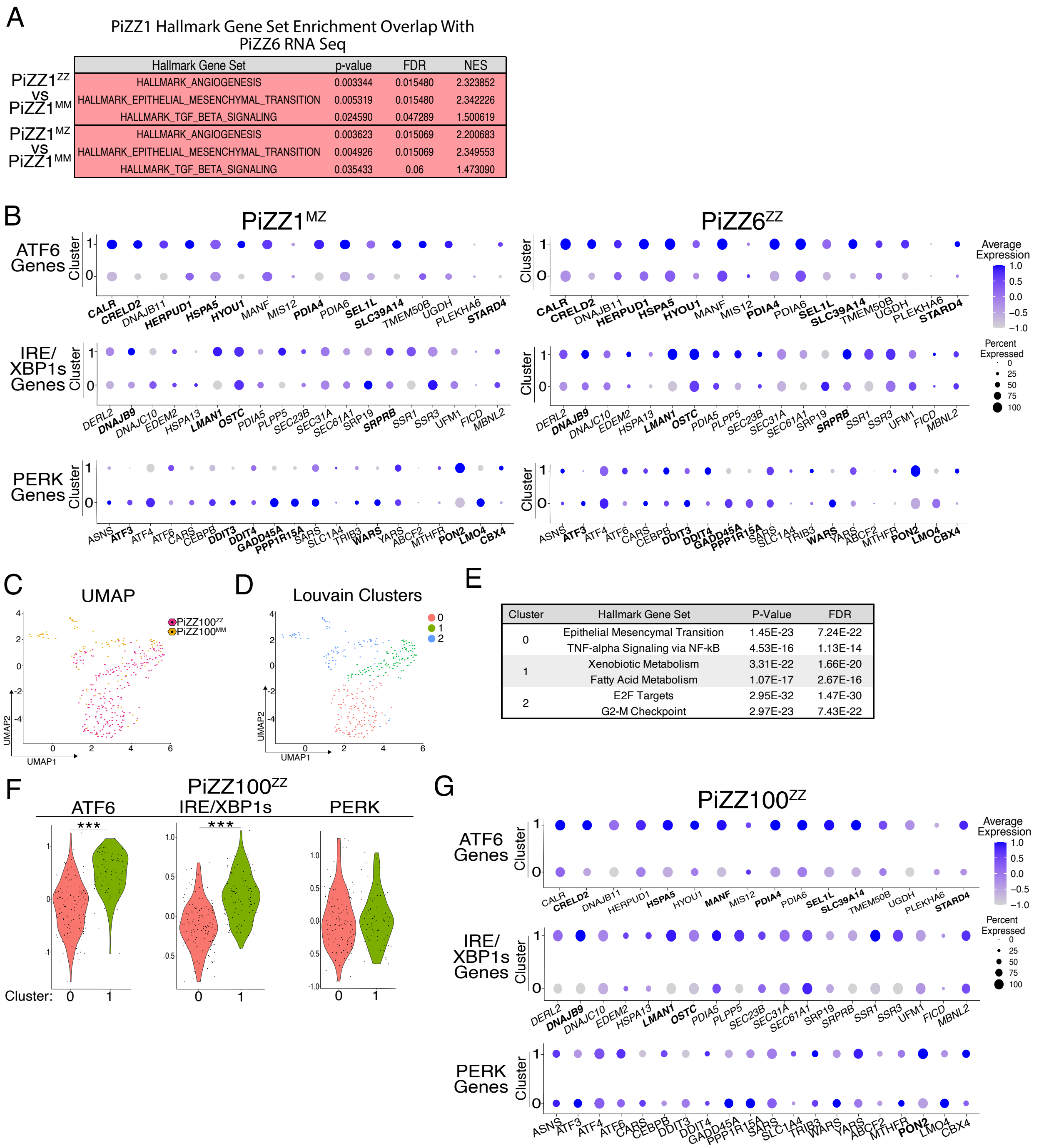


| | Hallmark Gene Set | p-value | FDR | NES |
|---|---|---|---|---|
| PiZZ1ZZ vs PiZZ1MM | HALLMARK_ANGIOGENESIS | 0.003344 | 0.015480 | 2.323852 |
| | HALLMARK_EPITHELIAL_MESENCHYMAL_TRANSITION | 0.005319 | 0.015480 | 2.342226 |
| | HALLMARK_TGF_BETA_SIGNALING | 0.024590 | 0.047289 | 1.500619 |
| PiZZ1MZ vs PiZZ1MM | HALLMARK_ANGIOGENESIS | 0.003623 | 0.015069 | 2.200683 |
| | HALLMARK_EPITHELIAL_MESENCHYMAL_TRANSITION | 0.004926 | 0.015069 | 2.349553 |
| | HALLMARK_TGF_BETA_SIGNALING | 0.035433 | 0.06 | 1.473090 |

| Cluster | Hallmark Gene Set | P-Value | FDR |
|---|---|---|---|
| 0 | Epithelial Mesencymal Transition | 1.45E-23 | 7.24E-22 |
| | TNF-alpha Signaling via NF-kB | 4.53E-16 | 1.13E-14 |
| 1 | Xenobiotic Metabolism | 3.31E-22 | 1.66E-20 |
| | Fatty Acid Metabolism | 1.07E-17 | 2.67E-16 |
| 2 | E2F Targets | 2.95E-32 | 1.47E-30 |
| | G2-M Checkpoint | 2.97E-23 | 7.43E-22 |

**Figure S3: Transcriptional Heterogeneity and Branch-Specific Activation of the UPR In MZ and ZZ iHeps**. A) GSEA analysis of PiZZ1 syngeneic iHeps grouped by original identity displaying overlapping Hallmark pathway enrichments from bulk RNA-seq (**Figure 2**). B) PiZZ1$^{MZ}$ and PiZZ6$^{ZZ}$ UPR branch specific gene dot plot projections. Genes that are differentially expressed are bolded (FDR p<0.05). C) UMAP projection by original identity for PiZZ100 ZZ and MM scRNA-seq experiment using the Fluidigm C1 platform. D) Louvain clustering demonstrates 3 clusters. E) Top two Hallmark gene sets by cluster as ranked by FDR using Enrichr analysis of all DEGs (FDR <0.05). F) Violin plots for UPR branch specific module scores for PiZZ100$^{ZZ}$ clusters 0 and 1. G) PiZZ100$^{ZZ}$ UPR branch specific gene dot plot projections. Genes that are differentially expressed are bolded (FDR p<0.05). ***p<0.001.

**Table S1: Top 20 GO terms enriched in MM iHeps compared with either MZ or ZZ iHeps as ranked by NES.**

| MZ vs MM | | | | ZZ vs MM | | | |
|---|---|---|---|---|---|---|---|
| GO TERM | p-value | FDR | NES | GO TERM | p-value | FDR | NES |
| GO_STEROL_HOMEOSTASIS | 0.00074 | 0.03346 | -2.39231 | GO_STEROL_HOMEOSTASIS | 0.00074 | 0.02275 | -2.45276 |
| GO_LIPID_HOMEOSTASIS | 0.00069 | 0.03346 | -2.27034 | GO_LIPID_HOMEOSTASIS | 0.00069 | 0.02275 | -2.37286 |
| GO_TRIGLYCERIDE_METABOLIC_PROCESS | 0.00073 | 0.03346 | -2.20907 | GO_PROTEIN_LIPID_COMPLEX_SUBUNIT_ORGANIZATION | 0.00080 | 0.02275 | -2.35827 |
| GO_PROTEIN_CONTAINING_COMPLEX_REMODELING | 0.00083 | 0.03346 | -2.18950 | GO_PROTEIN_CONTAINING_COMPLEX_REMODELING | 0.00082 | 0.02275 | -2.35273 |
| GO_PROTEIN_ACTIVATION_CASCADE | 0.00083 | 0.03346 | -2.18073 | GO_REGULATION_OF_STEROID_METABOLIC_PROCESS | 0.00070 | 0.02275 | -2.33170 |
| GO_PROTEIN_LIPID_COMPLEX_SUBUNIT_ORGANIZATION | 0.00081 | 0.03346 | -2.17964 | GO_STEROL_BIOSYNTHETIC_PROCESS | 0.00075 | 0.02275 | -2.32424 |
| GO_ORGANIC_HYDROXY_COMPOUND_CATABOLIC_PROCESS | 0.00076 | 0.03346 | -2.16887 | GO_STEROL_METABOLIC_PROCESS | 0.00067 | 0.02275 | -2.31082 |
| GO_BRUSH_BORDER_MEMBRANE | 0.00079 | 0.03346 | -2.16585 | GO_REGULATION_OF_PLASMA_LIPOPROTEIN_PARTICLE_LEVELS | 0.00074 | 0.02275 | -2.30332 |
| GO_STEROID_ESTERIFICATION | 0.00086 | 0.03346 | -2.15333 | GO_STEROL_TRANSPORT | 0.00074 | 0.02275 | -2.26510 |
| GO_CELLULAR_BIOGENIC_AMINE_METABOLIC_PROCESS | 0.00078 | 0.03346 | -2.14896 | GO_ORGANIC_ACID_CATABOLIC_PROCESS | 0.00063 | 0.02275 | -2.25546 |
| GO_DIGESTION | 0.00071 | 0.03346 | -2.11228 | GO_REGULATION_OF_CHOLESTEROL_METABOLIC_PROCESS | 0.00076 | 0.02275 | -2.24717 |
| GO_CHOLESTEROL_EFFLUX | 0.00080 | 0.03346 | -2.10765 | GO_STEROID_BIOSYNTHETIC_PROCESS | 0.00066 | 0.02275 | -2.24666 |
| GO_MICROVILLUS | 0.00075 | 0.03346 | -2.10161 | GO_MONOCARBOXYLIC_ACID_CATABOLIC_PROCESS | 0.00069 | 0.02275 | -2.22795 |
| GO_REGULATION_OF_STEROID_METABOLIC_PROCESS | 0.00071 | 0.03346 | -2.09505 | GO_BILE_ACID_METABOLIC_PROCESS | 0.00078 | 0.02275 | -2.22404 |
| GO_STEROL_TRANSPORTER_ACTIVITY | 0.00082 | 0.03346 | -2.09457 | GO_REGULATION_OF_TRIGLYCERIDE_METABOLIC_PROCESS | 0.00083 | 0.02275 | -2.21593 |
| GO_NEUTRAL_LIPID_BIOSYNTHETIC_PROCESS | 0.00080 | 0.03346 | -2.08586 | GO_PROTEIN_LIPID_COMPLEX_ASSEMBLY | 0.00082 | 0.02275 | -2.21311 |
| GO_STEROL_METABOLIC_PROCESS | 0.00068 | 0.03346 | -2.08177 | GO_STEROID_METABOLIC_PROCESS | 0.00062 | 0.02275 | -2.20904 |
| GO_MYELIN_ASSEMBLY | 0.00083 | 0.03346 | -2.07211 | GO_TRIGLYCERIDE_METABOLIC_PROCESS | 0.00074 | 0.02275 | -2.20839 |
| GO_NEUTRAL_LIPID_METABOLIC_PROCESS | 0.00071 | 0.03346 | -2.06520 | GO_REGULATION_OF_LIPID_BIOSYNTHETIC_PROCESS | 0.00066 | 0.02275 | -2.20832 |
| GO_STEROL_TRANSPORT | 0.00073 | 0.03346 | -2.04958 | GO_REGULATION_OF_LIPID_CATABOLIC_PROCESS | 0.00076 | 0.02275 | -2.19853 |

**Table S2: Differential Metabolites ranked by FDR**

| Metabolite | p-value | FDR |
|---|---|---|
| **Amide Metabolites** | | |
| Glycocholic acid | 3.11E-12 | 1.54E-10 |
| Kynurenine | 3.27E-12 | 1.54E-10 |
| Pantothenic acid | 4.71E-12 | 1.54E-10 |
| Glycerol-3-phosphate | 7.80E-12 | 1.91E-10 |
| N-Acetyl-L-Glutamic acid | 6.47E-10 | 1.27E-08 |
| Glutathione_reduced | 1.00E-09 | 1.63E-08 |
| UDP-GlcNAc | 1.29E-09 | 1.80E-08 |
| D-Gluconic acid | 1.50E-09 | 1.84E-08 |
| UDP-glucose / -galactose | 3.74E-09 | 4.07E-08 |
| Glycochenodeoxycholic acid | 9.84E-09 | 9.39E-08 |
| Hypoxanthine | 1.05E-08 | 9.39E-08 |
| Pyruvic acid | 1.42E-08 | 1.16E-07 |
| Pseudouridine | 1.58E-07 | 1.17E-06 |
| ATP | 1.79E-07 | 1.17E-06 |
| Glyceric acid | 3.46E-07 | 2.12E-06 |
| AMP | 5.10E-07 | 2.77E-06 |
| N-Acetyl-L-Methionine | 8.00E-07 | 4.12E-06 |
| Citrulline | 1.36E-06 | 6.68E-06 |
| Kynurenic acid | 2.18E-06 | 1.02E-05 |
| ADP | 3.31E-06 | 1.47E-05 |
| Uric acid | 3.69E-06 | 1.57E-05 |
| Oxypurinol | 6.97E-06 | 2.84E-05 |
| Xanthine | 1.05E-05 | 4.12E-05 |
| 3-Hydroxybutyric acid | 1.31E-05 | 4.94E-05 |
| Glutathione Disulfide | 2.58E-05 | 9.36E-05 |
| Taurodeoxycholic acid<br>Taurochenodeoxycholic acid | 3.69E-05 | 0.0001 |
| 2-ketoisovaleric acid KIV | 8.88E-05 | 0.0003 |
| Arachidonic acid | 0.0002 | 0.0005 |
| 2-Hydroxyglutaric acid | 0.0002 | 0.0005 |
| Uridine | 0.0002 | 0.0006 |
| cyclic-AMP | 0.0002 | 0.0006 |
| Oxalic acid | 0.0002 | 0.0006 |
| 1,5-AG / 1-deoxyglucose | 0.0002 | 0.0006 |
| Glucose/Fructose/Galactose_waterloss | 0.0003 | 0.0007 |
| a-Ketoglutaric acid | 0.0004 | 0.0010 |
| Indole-3-lactic acid | 0.0007 | 0.0019 |
| Acetoacetic acid | 0.0022 | 0.0056 |
| Butyric acid | 0.0024 | 0.0058 |
| Inosine | 0.0030 | 0.0072 |
| Anthranilic acid | 0.0039 | 0.0091 |
| Keto-isocaproic acid KIC<br>Keto-methylvalerate KMV | 0.0040 | 0.0091 |
| Methylmalonic acid | 0.0042 | 0.0093 |
| Taurine | 0.0056 | 0.0117 |
| Adipic acid | 0.0056 | 0.0117 |
| N-Acetyl-L-Aspartic acid | 0.0067 | 0.0138 |
| N-Acetyl-L-Glutamine | 0.0071 | 0.0142 |
| Oleoyl Glycine | 0.0096 | 0.0188 |
| Lactic acid | 0.0113 | 0.0214 |
| Glutamic acid | 0.0117 | 0.0216 |
| Succinic acid | 0.0126 | 0.0228 |
| Aconitic acid | 0.0138 | 0.0245 |
| Oleoyl Phenylalanine | 0.0190 | 0.0332 |
| Citric acid/Isocitric acid | 0.0213 | 0.0367 |
| Deoxycholic acid | 0.0233 | 0.0393 |
| Hippuric acid | 0.0251 | 0.0416 |

| Metabolite | p-value | FDR |
|---|---|---|
| **Lipid Metabolites** | | |
| kynurenine | 4.50E-16 | 5.36E-14 |
| proline | 3.46E-15 | 2.06E-13 |
| asparagine | 3.30E-12 | 1.31E-10 |
| serine | 7.31E-12 | 1.76E-10 |
| N-carbomoyl-beta-alanine | 8.84E-12 | 1.76E-10 |
| 5-Adenosyl-Methionine (SAMe) | 8.85E-12 | 1.76E-10 |
| alpha-glycerophosphocholine | 4.16E-11 | 6.61E-10 |
| citrulline | 4.59E-11 | 6.61E-10 |
| phosphoethanolamine | 5.49E-11 | 6.61E-10 |
| N-Acetyl-Lysine | 5.56E-11 | 6.61E-10 |
| kynurenic acid | 7.15E-11 | 7.74E-10 |
| Ala-Leu | 1.98E-10 | 1.94E-09 |
| citicholine | 2.12E-10 | 1.94E-09 |
| C9-carnitine | 6.37E-10 | 5.41E-09 |
| serotonin | 8.88E-10 | 7.04E-09 |
| betaine | 1.69E-09 | 1.17E-08 |
| 5-HIAA | 1.75E-09 | 1.17E-08 |
| creatine | 1.77E-09 | 1.17E-08 |
| N-Acetyl-L-Methionine | 2.27E-09 | 1.42E-08 |
| ADMA/SDMA | 4.20E-09 | 2.50E-08 |
| carnitine | 6.12E-09 | 3.47E-08 |
| C14-carnitine | 1.34E-08 | 7.25E-08 |
| N-Carbamoyl-BAIBA | 1.63E-08 | 8.13E-08 |
| C3-malonyl-carnitine | 1.64E-08 | 8.13E-08 |
| N1-Methylnicotinamide | 1.80E-08 | 8.55E-08 |
| acetylcholine | 2.67E-08 | 1.22E-07 |
| anthranilic acid | 5.14E-08 | 2.27E-07 |
| C4-methylmalonyl-carnitine | 7.58E-08 | 3.22E-07 |
| C4-butyryl-carnitines | 8.89E-08 | 3.65E-07 |
| C5-valeryl-carnitines | 1.50E-07 | 5.96E-07 |
| C3-carnitine | 2.32E-07 | 8.91E-07 |
| Cystine | 2.58E-07 | 9.61E-07 |
| N-Acetyl-L-Phenylalanine | 2.76E-07 | 9.97E-07 |
| N-Acetyl-L-Alanine | 3.44E-07 | 1.21E-06 |
| C16-carnitine | 1.13E-06 | 3.84E-06 |
| creatinine | 1.23E-06 | 4.08E-06 |
| glycine | 1.48E-06 | 4.75E-06 |
| glutamate | 1.69E-06 | 5.29E-06 |
| valine | 1.83E-06 | 5.57E-06 |
| NMMA | 2.14E-06 | 6.37E-06 |
| C6-carnitine | 3.29E-06 | 9.48E-06 |
| threonine | 3.34E-06 | 9.48E-06 |
| Norepinephrine-updated | 6.14E-06 | 1.69E-05 |
| C2-carnitine | 6.29E-06 | 1.69E-05 |
| histidine | 6.41E-06 | 1.69E-05 |
| C18-carnitine | 8.97E-06 | 2.32E-05 |
| aminoisobutyric acid | 1.11E-05 | 2.81E-05 |
| 1-methylhistamine | 1.24E-05 | 3.06E-05 |
| N-Methyl-4-Pyridone-3-Carboxamide | 1.68E-05 | 4.08E-05 |
| cis/trans hydroxyproline | 2.10E-05 | 5.00E-05 |
| N-Acetyl-L-Ornithine | 2.36E-05 | 5.52E-05 |
| C18:1-carnitine | 2.79E-05 | 6.38E-05 |
| 2'-deoxycytidine | 3.08E-05 | 6.91E-05 |
| arginine | 3.14E-05 | 6.91E-05 |
| 2'-deoxyadenosine | 5.03E-05 | 0.0001 |
| uridine | 6.11E-05 | 0.0001 |
| N-Acetyl-L-Glutamic acid | 6.74E-05 | 0.0001 |
| C8-carnitine | 8.40E-05 | 0.0002 |
| 3-hydroxyanthranilic acid | 8.95E-05 | 0.0002 |
| 5'-adenosylhomocysteine | 9.78E-05 | 0.0002 |
| Dihydrouracil T1 | 0.0001 | 0.0002 |
| carnosine | 0.0001 | 0.0002 |
| tryptophan | 0.0001 | 0.0003 |
| taurine | 0.0002 | 0.0003 |
| glutamine | 0.0002 | 0.0003 |
| aspartate | 0.0002 | 0.0003 |
| phenylalanine | 0.0003 | 0.0005 |
| histamine | 0.0003 | 0.0005 |
| lysine | 0.0004 | 0.0007 |
| GABA | 0.0005 | 0.0008 |
| glycerol | 0.0005 | 0.0009 |
| C12-carnitine | 0.0008 | 0.0014 |
| dimethylglycine | 0.0011 | 0.0018 |
| leucine | 0.0012 | 0.0020 |
| xanthurenate | 0.0014 | 0.0023 |
| ornithine | 0.0015 | 0.0024 |
| choline | 0.0016 | 0.0025 |
| Ala-Gly | 0.0017 | 0.0027 |
| methyl-hydroxyisobutyric acid | 0.0018 | 0.0027 |
| methionine | 0.0021 | 0.0031 |
| DMGV T1 | 0.0030 | 0.0043 |
| cAMP | 0.0044 | 0.0064 |
| thiamine | 0.0071 | 0.0101 |
| spermidine | 0.0099 | 0.0141 |
| C5-glutaryl-carnitine | 0.0110 | 0.0154 |
| niacinamide | 0.0112 | 0.0155 |
| tyrosine | 0.0129 | 0.0176 |
| alanine | 0.0146 | 0.0197 |
| Sarcosine | 0.0158 | 0.0211 |
| thymidine | 0.0166 | 0.0220 |
| C7-carnitine | 0.0199 | 0.0260 |
| isoleucine | 0.0228 | 0.0292 |
| N-Methyl-2-Pyridone-5-Carboxamide | 0.0228 | 0.0292 |
| phosphocholine | 0.0236 | 0.0299 |
| Dopamine-updated | 0.0268 | 0.0336 |
